# Safeguarding epithelial junctions by a novel quality control pathway

**DOI:** 10.64898/2026.08.11.744142

**Authors:** Cristina Tocchini, Angelo L. Angonezi, Dania Camila Pulido Barrera, Susan E. Mango

## Abstract

Cell junctions establish and maintain epithelial architecture despite fluctuating environmental and developmental conditions. A central question is how cells respond to challenging conditions to preserve junctional integrity. Here, we report the discovery of a previously unrecognized quality control pathway that monitors epithelial junctions (J-QC). We used the *Caenorhabditis elegans* epidermis as a model to investigate the DLG-1-AJM-1 complex (DAC), a junctional domain that is critical for embryonic morphogenesis. We identify two mechanisms that sustain junctional integrity: first, localized *dlg-1* mRNA ensures appropriate DLG-1 protein levels at the junction; repositioning *dlg-1* RNA reduces DLG-1 levels, leading to gaps between epithelial cells. Second, transcription of DAC components responds to perturbations that disrupt the DAC. This response is sequence-independent, distinguishing it from other quality control mechanisms. It is activated by perturbations of the DAC or cytoskeleton and requires the LINC complex component ZYG-12/HOOK1-3 to transduce information about junctional integrity to the nucleus. These findings define a novel junctional QC for epithelial maintenance.

---

Epithelial junctions, such as adherens and tight junctions, organize and compartmentalize tissues^1–3^. By forming continuous contacts between neighboring cells, epithelial junctions establish an apical-basolateral boundary, maintain epithelial cohesion, and create barriers that separate the internal from the external environment^4^. However, little is known about how established junctions maintain homeostasis as epithelia experience developmental remodeling and mechanical stress. This question carries broad biological and biomedical significance, as disruption of epithelial junctions underlies pathological conditions, including the epithelial-to-mesenchymal transition during cancer progression and intestinal barrier dysfunction during aging^5–7^. Junctional integrity is equally critical during embryogenesis, where junctions contribute to morphogenesis and body axis elongation in many organisms^2,8–10^. During these processes, epithelial junctions are subjected to persistent mechanical and developmental stimuli, including cellular rearrangements and body elongation, which continuously challenge their integrity. Extrusion of damaged cells is one established route for preserving epithelial integrity, executed non-autonomously by neighboring cells^11^. Mechanisms that maintain epithelia by a cell-autonomous route are largely unknown.

Here, we uncover how *C. elegans* embryonic epidermal cells maintain cell junction integrity during morphogenesis. Using a genetic approach combined with imaging and quantitation analyses, we find that RNA localization sustains appropriate protein levels of the junction component DLG-1 and maintains DAC architecture. In addition, we identify a novel Junctional Quality Control (J-QC) pathway that surveils the DAC and, upon perturbation, upregulates transcription of DAC components. The J-QC relies on cytoskeletal and LINC components to relay information from the junction to the nucleus. Unlike previously described QC pathways, which monitor organelle integrity^12^ or the fidelity of molecular processes^13^, the J-QC surveils a subcellular structure, the junction, and couples its disruption to a compensatory transcriptional response. Together, these define a mode of cellular surveillance dedicated to maintaining a specific architectural structure within the cell.

## Results

### Subcellular localization of *dlg-1* mRNA is required for proper DLG-1 protein levels at epidermal junctions

*C. elegans*, epithelial junctions share structural, molecular, and regulatory features of adherens and tight junctions present in other organisms, reflecting broad evolutionary conservation of junctional architecture and its regulatory logic^14–16^. The junctional components in *C. elegans* are organized into two domains, the cadherin-catenin complex (CCC) and the DLG-1-AJM-1 complex (DAC)^17–19^, which are critical for barrier formation and tissue morphogenesis. Loss of any of these components leads to morphogenetic defects^20^, caused by the rupture of junctions under mechanical tension during embryo elongation, and ultimately death^21^.

Multiple transcripts coding for components in epithelial junctions preferentially localize to the junction itself during *Caenorhabditis elegans* morphogenesis^22^. To determine the roles of this subcellular localization, we redirected *dlg-1* mRNA away from the junction using our recently established MS2-MCP technique for *C. elegans* endogenous mRNAs^23^. As recent studies have shown that translation and protein folding can normally occur at the cytoplasmic side of the nuclear pore^24,25^, we decided to modify our MS2-MCP system to re-localized *dlg-1* mRNA at this location. Briefly, we fused the MS2 coat protein (MCP) to the nucleoporin NPP-9/Nup358 (NPP(+)) to generate an MCP::NPP-9 fusion protein (NPP(+)) as well as an unfused control (NPP(-)) Fig. 1A and S1A-F). Because NPP-9 localizes to the cytoplasmic filaments of the nuclear pore complex^26,27^, this fusion protein tethered *dlg-1* mRNAs carrying MS2 hairpins (MS2(+)) to the cytoplasmic face of the NPC (Fig. 1A,B). Consistent with this idea, perinuclear *dlg-1* mRNAs colocalized with MCP::NPP-9 (Fig. 1B). This approach redirected endogenous *dlg-1* transcripts away from the junction while keeping the mRNA configuration identical between test and control conditions.

**Figure 1.**
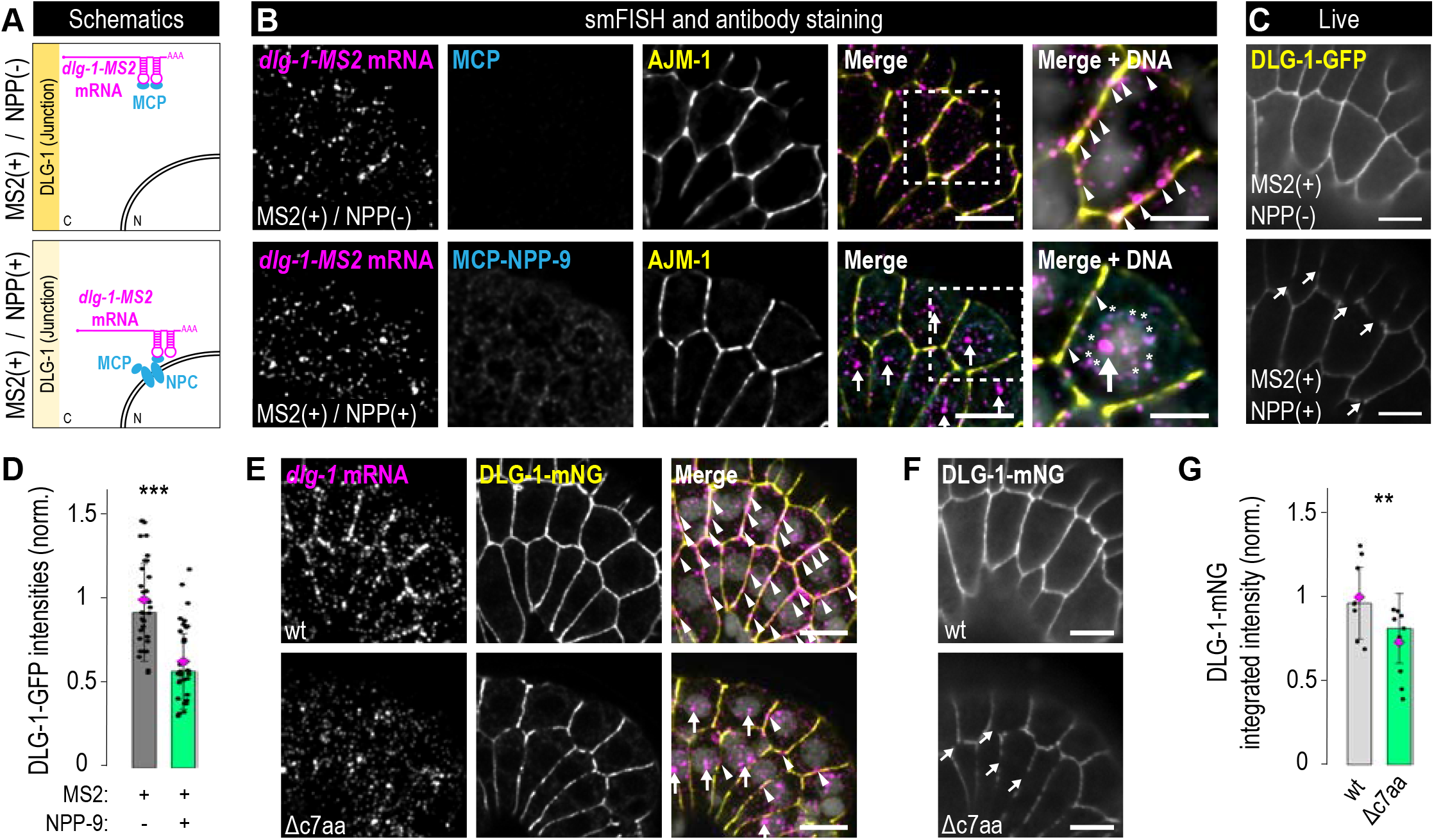
Localization of *dlg-1* transcripts controls junctional DLG-1 protein levels. **A.** Schematic representation of an epidermal cell with the adherens junction depicted in yellow as described in Fig. S1A. *dlg-1* mRNA with MS2 hairpins in its 3’UTR (MS2(+), magenta). In cyan, upper panel: MCP alone (NPP(-)) bound to *dlg-1MS2* mRNA at junctions; lower panel: MCP fused to NPP-9/NUP358 (NPP(+)), bound to *dlg-1MS2* mRNA at nuclear pores. **B.** Fluorescence images of epidermal cells of smFISH for *dlg-1* mRNA (magenta), MCP (upper) or MCP::NPP-9 (lower) (cyan), and junctional AJM-1 (yellow). Merged images and zoom-ins of one cell (dotted square) are shown. Transcripts (magenta, arrowheads) localized at the junction (yellow). mRNAs (magenta) colocalize with MCP::NPP-9 perinuclearly (asterisks). Vertical arrows depict large foci within nuclei (transcription sites). Scale bars: 5 µm and 2.5 µm (zoom-ins). **C.** Live fluorescence images of epidermal cells show gaps (arrows) in junctions (DLG-1::GFP) for cells with re-localized *dlg-1* RNA (lower) compared to controls (upper). Scale bars: 5 µm. **D.** Bar plot with error bars normalized to the control MS2(+); NPP(-) (dark grey) and showing the reduction of DLG-1::GFP fluorescence intensities in MS2(+); NPP(+) (green) strains, where *dlg-1* transcripts are re-localized to nuclear pores. Each dot represents the maximum fluorescent intensity of a junction shared between two seam cells (three junctions from the posterior-most seam cells per embryo) marked with endogenous DLG-1::GFP. *** = p < 0.001. For raw data and statistics, see Table S3. **E.** Fluorescence images of epidermal cells of smFISH for *dlg-1* mRNA (magenta), DLG-1::mNG protein (yellow), and merge with DNA (white). Upper panels: wild type. Lower panels: *dlg-1(Δc7aa).* Arrowheads depict *dlg-1* transcripts (magenta) colocalizing with DLG-1 protein at the junction (yellow). Vertical arrows depict large foci within the nuclei (transcription sites). Scale bars: 5 µm. **F.** Live fluorescence images of epithelia for wild-type DLG-1(wt) (upper) and DLG-1(Δc7aa) (lower). Arrows depict gaps in the junctional DAC (DLG-1::mNG). Scale bars: 5 µm. **G.** Bar plot with error bars showing the median (magenta) and SD (black) normalized to the wild-type control (light grey). Note the reduction in DLG-1(Δc7aa)::mNG fluorescence (green). Each dot represents the mean integrated fluorescent intensity of endogenous DLG-1::mNG signal from a single Z-stack of whole embryos focusing on epidermal cells. For raw data and statistics, see Table S3.

Quantitation of junctional fluorescent intensities^23^ revealed a 50% reduction of DLG-1::GFP fluorescence at the junctions of MS2(+) strains with mRNA localized to nuclear pores (NPP(+)) compared to controls (NPP(-)) (Fig. 1C,D). The variability observed in DLG-1 levels among both test and control strains likely reflects subtle embryo-to-embryo differences, but levels were clearly lower following re-localization. Normally, DLG-1 protein levels rise as embryogenesis progresses. We observed the same trend in the re-localization strains, suggesting this aspect of *dlg-1* regulation remained intact, but DLG-1 never achieved the levels seen in wild-type embryos (data not shown). As expected, when *dlg-1* mRNA did not contain MS2 hairpins (MS2(-)), transcripts localized normally at junctions in both NPP(-)^23^ and NPP(+) backgrounds (Fig. S1B,G,H). In this configuration, DLG-1::GFP protein exhibited comparable fluorescence levels in NPP(-) and NPP(+) conditions, (Fig. S1I,J). This result indicates that the reduced DLG-1 levels observed upon transcript re-localization were not due to secondary effects caused by transgenic NPP-9 expression. We conclude that localized mRNA is critical to establish normal levels of DLG-1 protein at junctions.

In strains with re-localized mRNA (MS2(+); NPP(+) configuration), but not in controls, we observed gaps along the circumference of the DLG-1-marked junctions (Fig. 1C). These gaps are consistent with impaired translation or distribution of wild-type DLG-1 protein resulting from re-localization of its mRNA. Gaps were also observed for AJM-1, a second component of the DAC, suggesting biogenesis of the DAC was delayed or disturbed (Fig. 1B). These data reveal the importance of mRNA subcellular localization to maintain appropriate protein levels and distribution along the periphery of the cell, to ensure junctional integrity.

To complement the mRNA re-localization approach, we sought to perturb junctional *dlg-1* mRNA localization through an independent genetic strategy. Our previous work found that the carboxy-terminal part of DLG-1 is required for mRNA localization but dispensable for protein localization in transgenic strains^22^. Based on these findings, we generated a *dlg-1* CRISPR allele lacking seven conserved amino acids within the sequence connecting SH3 and GuK domains (*dlg-1(Δc7aa)*; Fig. S2A) in a strain carrying an mNeonGreen (mNG) fluorescent tag at the carboxy-terminus of DLG-1 (LP598^50^). Previous work demonstrated that the mNG tagged DLG-1 is fully functional and appropriately localized in the wild-type configuration^28^. Single molecule fluorescent *in situ* hybridization (smFISH) analyses revealed that endogenous *dlg-1* mRNAs carrying the Δc7aa mutation failed to localize properly to embryonic epithelial junctions compared to wild-type controls (Fig. 1E). To determine the consequences of disrupting *dlg-1* mRNA localization through this endogenous mutation, we analyzed junction morphology by live imaging. Similar to the RNA re-localization experiment, *dlg-1(Δc7aa)* embryos displayed discontinuities in junctional organization, characterized by the appearance of gaps along epithelial junctions (Fig. 1E,F and S2B). Similarly to what we observed in our re-localization experiments, quantitation of DLG-1 fluorescence intensities revealed a reduction in mean integrated intensity of junctional DLG-1(Δc7aa) signal relative to controls (Fig. 1G). The reduction was less dramatic than the nuclear pore experiment, possibly because some transcripts were still localized at the junction (Fig. 1E), but the reduction was statistically significant. These experiments show that *dlg-1* mRNA localization is required for appropriate DLG-1 protein levels, and re-localization disrupts junctional integrity. Thus, mRNA localization contributes to the robustness of epithelial junctions during morphogenesis.

### Quantitation of transcriptional levels by DotQuant

We wondered why animals survived despite mis-localized *dlg-1* mRNA, reduced protein levels, and junctional gapping. Close inspection revealed that there was an increase in *dlg-1* transcription when transcripts were re-localized. For example, smFISH experiments of *dlg-1* mRNA re-localized by MS2 and MCP::NPP-9 had larger and brighter foci in epidermal nuclei of MS2(+); NPP(+) strains compared to controls (Fig. 1B,E). These dots represented transcription sites, based on smFISH for intronic sequences, and their increased size and brightness suggested an upregulation in transcription^29^. To quantify transcriptional changes in nuclei from different conditions, we developed a rapid pipeline for nuclear dot quantitation (DotQuant, https://github.com/angelo-angonezi/DotQuant). DotQuant combines automated nuclear and three-dimensional spot segmentation and fluorescence measurements in a single workflow and is supported by custom computational tools that enable fast and reproducible analysis of transcription sites across large numbers of nuclei. The pipeline provides the integrated intensity (volume × mean signal intensity) of individual transcription sites as a quantitative measure of transcriptional activity (Fig. S3A-J; Methods). DotQuant enabled quantitative comparison of transcriptional output between different genetic conditions, providing a robust approach to assess changes in *dlg-1* transcriptional activity in our study.

### Re-localized *dlg-1* mRNA triggers compensatory upregulation of *dlg-1* transcription, which supports viability

Quantitation of the integrated intensity of these transcription sites by DotQuant showed a 3.4-fold increase for re-localized *dlg-1* transcripts compared to controls (Fig. 2A and S3A-J). We conclude that cells upregulate *dlg-1* transcription when *dlg-1* mRNAs are mis-localized.

**Figure 2.**
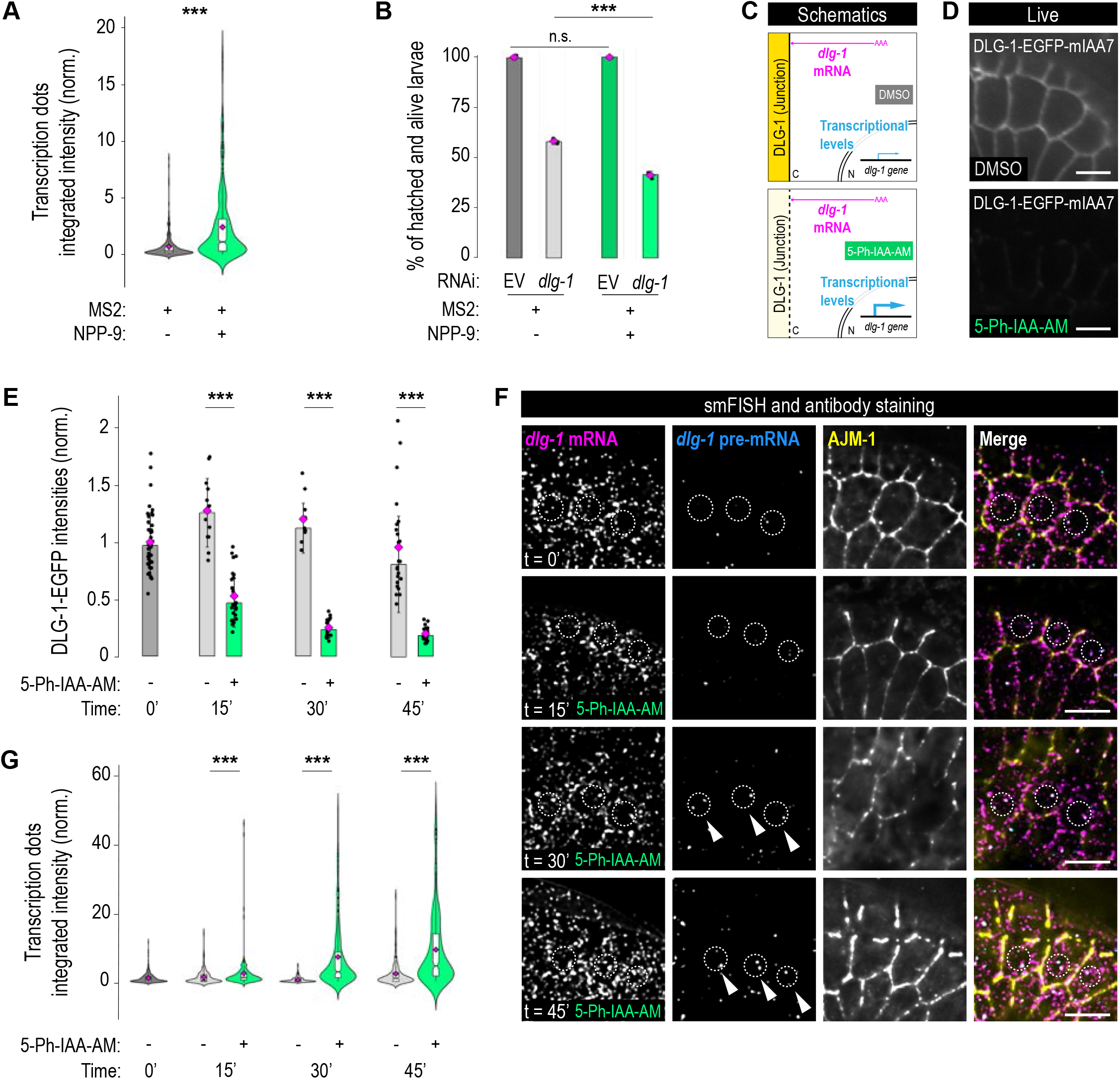
Depletion of DLG-1 protein induces *dlg-1* gene transcriptional. **A.** Intensity of nuclear mRNA dots increases after RNA re-localization. Violin plots with overlaid notched boxplots of distributions of integrated intensities of transcription dots across strains. *** = p < 0.001. For raw data and statistics, see Table S4. **B.** Bar plot with error bars for embryonic viability in re-localized *dlg-1* (light green NPP(+)) versus control (grey) after *dlg-1* RNAi. Each dot represents the percentage of hatched and living larvae per biological replicate (x3). n.s. = p > 0.05; *** = p < 0.001. For raw data and statistics, see Table S3. **C.** Schematic representation of a portion of a seam cell as described in Fig. S1A. Upper panel: wild-type levels of DLG-1 protein at the junction (yellow); wild-type transcriptional levels (thin arrow) of *dlg-1* gene (cyan); addition of DMSO (grey) as a negative control. Lower panel: reduced levels of DLG-1 protein at the junction (light yellow); increased transcriptional levels (thicker arrow) of *dlg-1* gene (cyan); addition of 5-Ph-IAA (green) as treatment to deplete DLG-1 protein with the AID2 system. In both instances, wild-type localization of *dlg-1* transcripts at the junction (magenta). **D.** Live fluorescence images of epidermal cells of fluorescence signal of DLG-1::EGFP::mIAA7 strain. In upper image: treatment with DMSO at t30 (control); lower image: treatment with 5-Ph-IAA-AM at t30 (green). Scale bars: 5 µm. **E.** Bar plot with error bars normalized to the reference t0 (dark grey) and showing the changes in DLG-1::EGFP fluorescence intensities throughout the time course experiment (light grey (-): DMSO treatment; green (+): 5-Ph-IAA-AM treatment). Each dot represents the maximum fluorescent intensity of a junction shared between two seam cells (three junctions from the posterior-most seam cells per embryo) marked with endogenous DLG-1::EGFP. ***: p < 0.001. For raw data and statistics, see Table S3. **F.** Fluorescence images of epidermal cells of smFISH signal for *dlg-1* mRNA (magenta) and *dlg-1* pre-mRNA (cyan), antibody staining of AJM-1 (yellow), and merge. From top to bottom, the time course experiments: immediately after placing embryos in M9 with DMSO (t0’); after 15 minutes in M9 with 5-Ph-IAA-AM (t15’); after 30 minutes in M9 with 5-Ph-IAA-AM (t30’); after 45 minutes in M9 with 5-Ph-IAA-AM (t45’). Example seam cell nuclei are circled with dashed lines. Arrowheads point at nuclei with larger transcription dots, found in later time points of the 5-Ph-IAA-AM treatment. Scale bar: 5 µm. **G.** Violin plots with overlaid notched boxplots of distributions of integrated intensities of transcription dots (channel for *dlg-1* pre-mRNA, cyan) in the time course experiment shown in (E). For raw data and statistics, see Table S4. **\*\*\*** = p < 0.001

We asked whether increased *dlg-1* transcription buffered the phenotypic consequences of RNA re-localization. We used partial RNAi (1:10 dilution) to reduce *dlg-1* mRNA levels, bringing the abundance of re-localized MS2(+); NPP(+) transcripts in line with that of localized MS2(+); NPP(-) control transcripts. Reduction of re-localized *dlg-1* mRNAs reduced viability to 40% whereas untreated strains of both genotypes were completely viable (98%; Fig 2B). This result shows that the increase in *dlg-1* transcripts promotes viability when mRNAs are mis-localized.

As a second test, we analyzed the *dlg-1(Δc7aa)* mutant, which also exhibits loss of *dlg-1* mRNA localization, reduced DLG-1 protein levels, and increased *dlg-1* transcription (Fig. 1E-G and S4A). We reasoned that if transcriptional upregulation contributes to survival of mutant DLG-1(Δc7aa), then partial reduction of *dlg-1(Δc7aa)* mRNA levels should compromise embryonic viability. Under control RNAi conditions, *dlg-1(Δc7aa)* embryos exhibited 80% embryonic viability compared to 99% in wild-type animals (Fig. S4B), suggesting that the conserved seven amino acids we deleted are critical for DLG-1. Upon mild *dlg-1* RNAi treatment, viability was reduced to 18% in *dlg-1(Δc7aa)* embryos compared to 53% in controls (Fig. S4B). Thus, partial reduction of *dlg-1* mRNA levels exacerbates the *dlg-1(Δc7aa)* mutant phenotype, supporting the conclusion that increased *dlg-1* transcription aids embryonic viability when junctional *dlg-1* function is compromised. Together, the data show that increased *dlg-1* transcription is functionally important and contributes to embryonic viability when *dlg-1* mRNA is mis-localized. These experiments demonstrate that augmented mRNA levels contribute to survival in cases where the DAC is challenged by *dlg-1* mutations or re-localization. We call this transcriptional response the junctional quality control pathway or J-QC, based on these and the next experiments.

### Reduction in DLG-1 protein triggers the *dlg-1* transcriptional increase

We hypothesized that increased *dlg-1* transcription responded either to mRNA re-localization *per se*, or to the consequent reduction in DLG-1 protein at the junction. To distinguish between these possibilities, we degraded DLG-1 protein while leaving *dlg-1* mRNA intact and localized appropriately (Fig. 2C). DLG-1 tagged with both mIAA7 and EGFP enabled us to induce and visualize DLG-1 degradation upon addition of the auxin analog 5-Ph-IAA-AM^30,31^ (Fig. 2C,D and S5A). In a time-course experiment, DLG-1::EGFP::mIAA7 fluorescence at the junction decreased by 50% within 15 minutes of treatment and 80% within 30-45 minutes, compared to time 0 (Fig. 2D,E). smFISH revealed that despite reduction of DLG-1 protein, junctional *dlg-1* mRNA localization was maintained (Fig. 2F, magenta and merge).

*dlg-1* transcription increased up to 15-fold after 45 minutes of treatment compared to controls (Fig. 2F,G and S5B). Whereas control cells had no or one transcriptional dot within cells, we observed many nuclei with two active loci after DLG-1 degradation, and these appeared within 30 minutes of treatment. Individual transcription dots were also brighter, compared to controls, suggesting that *dlg-1* loci were more active after DLG-1 degradation compared to controls. These findings support the idea that epithelial cells monitor DLG-1 protein levels and respond with a compensatory increase in *dlg-1* transcription to restore junctional protein abundance. Degradation of DLG-1 protein for 45 minutes or longer compromised the overall structure of the junction. Junctions no longer formed a continuous belt around the cell but instead appeared discontinuous, as visualized by AJM-1 (Fig. 2F, yellow, and S5C). This phenotype resembled the phenotypes observed after mRNA re-localization as well as published *dlg-1* loss-of-function contexts^32^ (RNAi or hypomorphic mutants). These data show that DLG-1 protein, not junctional *dlg-1* mRNA, mediates the J-QC pathway.

### A junctional quality control (J-QC) pathway selectively monitors DAC integrity

Some QC systems monitor via sequence complementarity (*e.g.,* transcriptional adaptation^33–35)^ whereas others are dedicated to a specific membranous organelle^12,13,36^. We asked whether the transcriptional upregulation of *dlg-1* reflected surveillance of a structure, namely the epithelial junction, or was specific for the *dlg-1* sequence. To distinguish between these possibilities, we analyzed a second component of the DAC called AJM-1 (Fig. 3A); *dlg-1* and *ajm-1* lack sequence similarity at the RNA or protein level. smFISH experiments coupled with quantitation of *dlg-1* transcription sites revealed significant upregulation of *dlg-1* in *ajm-1* mutants compared to a wild-type control (Fig. 3B,C). Conversely, 5-Ph-IAA-AM-mediated depletion of DLG-1 protein or RNAi-mediated inactivation of *dlg-1* transcripts led to a significant upregulation of *ajm-1* transcription (Fig 3D,E). We observed upregulation in developing epidermal cells, particularly in dorsal epidermal cells (Fig. 3D,E and S6A,B). Because *dlg-1* and *ajm-1* share no primary sequence similarity, these results indicate that the transcriptional response reflects the compromised integrity of the DAC epithelial junction rather than sequence complementarity. As predicted by this model, loss of *ajm-1* did not show additive effects upon simultaneous DLG-1 depletion, consistent with both perturbations acting in the same pathway. We refer to this pathway as the junctional quality control (J-QC), to distinguish it from previously described processes. We note that the nonsense-mediated mRNA decay pathway is required for transcriptional adaptation, but was dispensable for the J-QC, indicating that these are distinct transcriptional pathways (Fig. S6C).

**Figure 3.**
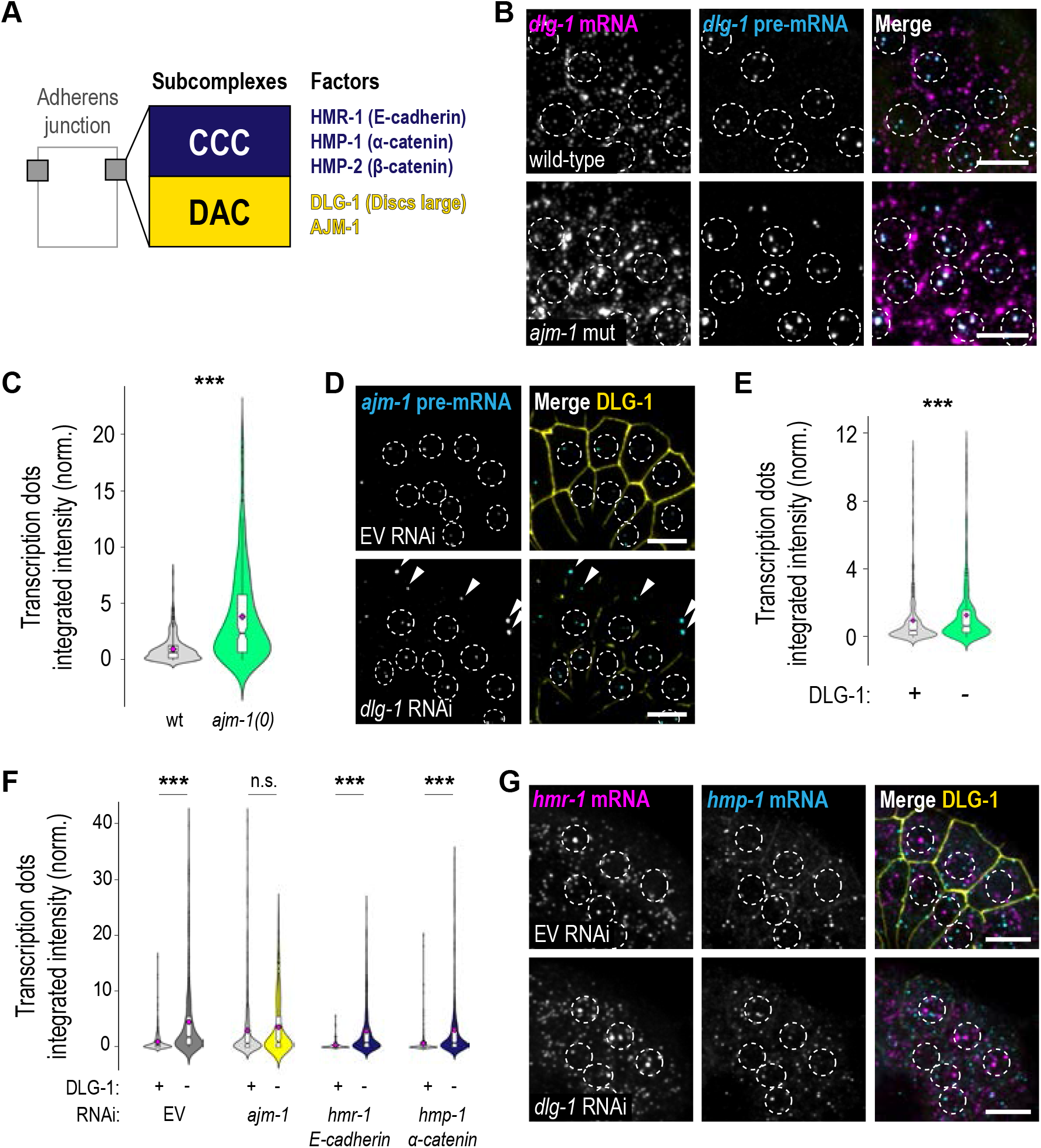
The DAC is a quality control checkpoint for the junctions. **A.** Simplified schematic representation of the two sub-complexes listing their components constituting *C. elegans* adherens junctions. An apical cadherin-catenin complex (CCC) in blue with HMR-1/E-cadherin, HMP-1/α-catenin, and HMP-2/β-catenin. Below the CCC, the DLG-1-AJM-1 complex (DAC) in yellow, named after its components. **B.** Fluorescence images of epidermal cells of smFISH signal for *dlg-1* mRNA (magenta) and *dlg-1* pre-mRNA (cyan), and merge. Upper panel: wild-type background. Lower panels: *ajm-1* mutant background. Example nuclei of seam and ventral epidermal cells are circled with dashed lines. Scale bar: 5 µm. **C.** Violin plots with overlaid notched boxplots of distributions of integrated intensities of transcription dots (channel for *dlg-1* pre-mRNA) in wild type versus *ajm-1* mutant backgrounds shown in (B). *** = p < 0.001. For raw data and statistics, see Table S4. **D.** Fluorescence images of epidermal cells of smFISH signal for *ajm-1* pre-mRNA (cyan), and DLG-1::EGFP (yellow). Upper panel: EV RNAi. Lower panels: *dlg-1* RNAi. Example nuclei of seam and ventral epidermal cells are circled with dashed lines. Arrowheads show transcription dots of dorsal epidermal cells. Scale bars: 5 µm. **E.** Violin plots with overlaid notched boxplots of distributions of integrated intensities of transcription dots for *ajm-1* pre-mRNAs in 5-Ph-IAA-AM-untreated (DLG-1 +) versus treated (DLG-1 -) DLG-1::EGFP::mIAA7 embryos. **\*\*\*** = p < 0.001. For raw data and statistics, see Table S4. **F.** Violin plots with overlaid notched boxplots of distributions of integrated intensities of transcription dots in multiple conditions. n.s. = p > 0.05; **\*\*\*** = p < 0.001. For raw data and statistics, see Table S4. **G.** Fluorescence images of epidermal cells of smFISH signal for *hmr-1* mRNA (magenta) and *hmp-1* mRNA (cyan), and merges with DLG-1::EGFP (yellow). Upper panel: EV RNAi. Lower panels: *dlg-1* RNAi. Example nuclei of seam and ventral epidermal cells are circled with dashed lines. Scale bars: 5 µm.

We next tested whether perturbation of alternative domains within epithelial junctions could trigger the J-QC. The cadherin-catenin complex (CCC) localizes apically to the DAC and is a separate compartment from the DAC (Fig. 3A). RNAi-mediated reduction of CCC components *hmr-1/E-cadherin* or *hmp-1/α-catenin* did not induce transcriptional upregulation of *dlg-1*, in contrast to *ajm-1* (Fig. 3F and S7A-C). Thus, disruption of the CCC alone was insufficient to activate the J-QC.

Next, we asked whether the CCC was required for J-QC triggered by DAC disruption. To test this idea, we combined *hmr-1* or *hmp-1* RNAi with AID2-mediated depletion of DLG-1 protein (Fig. S7B,C). Reduction of *hmr-1* or *hmp-1* did not alter *dlg-1* transcriptional upregulation compared to DLG-1 depletion alone (Fig. 3F and S6A). Thus, CCC components are not necessary for the J-QC response. Conversely, downregulation of *dlg-1* mRNA did not induce transcriptional upregulation of genes encoding CCC components or apical polarity regulators (*par-6* and *pkc-3*) (Fig. 3G and S7D). RT-qPCR analysis of intronic and exonic sequences in wild type and *ajm-1* mutant embryos confirmed selective upregulation of *dlg-1*, but not CCC components, in response to *ajm-1* disruption (Fig. S7E). These results indicate that the CCC and DAC, although adjacent within epithelial junctions, behave autonomously and without detectable cross-regulation. Thus, general cellular disturbances cannot account for *dlg-1* transcriptional induction, but rather, the effect is specific. In sum, the DAC employs a selective J-QC mechanism that responds precisely to perturbations affecting DAC integrity.

### The cytoskeleton acts as a bridge between the DAC and transcriptional regulation

We wondered how aberrant junctions signal to nuclei to alter gene expression. Given the involvement of junctions in cytoskeletal organization, we hypothesized that the cytoskeleton would be required to relay information from the junction to the nucleus. To minimize pleiotropic effects of cytoskeletal disruption, we took advantage of mutants with mild or inducible alterations in cytoskeletal organization. Specifically, we used a viable loss-of-function triple mutant to disrupt actin (*act-1,2,3(lf)*), and two thermo-sensitive mutants to disrupt the microtubule-based cytoskeleton: one for microtubules (*tba-1(ts)*), and one for dynein heavy chain (*dhc-1(ts)*), which drives minus-end-directed transport along microtubules. All three mutants showed a significant increase in *dlg-1* transcription compared to controls after a brief inactivation (two hours; Fig. 4A,B and S8A). *dhc-1(ts)* transcriptional levels were the highest of the three mutants, suggesting a pivotal role for the cytoskeleton in general, and microtubule-mediated movement in particular, for junctional homeostasis (Fig. 4A,B and S8A). We also found that auxin-mediated removal of DLG-1 protein showed no significant difference in *dlg-1* transcriptional signal when the cytoskeleton was disrupted (Fig. 4C,D and S8B). The lack of an additive effect upon simultaneous DLG-1 depletion and cytoskeletal disruption suggests that cytoskeletal components, microtubule-dependent retrograde transport, and DLG-1 function within the same pathway to mediate J-QC transcription.

**Figure 4.**
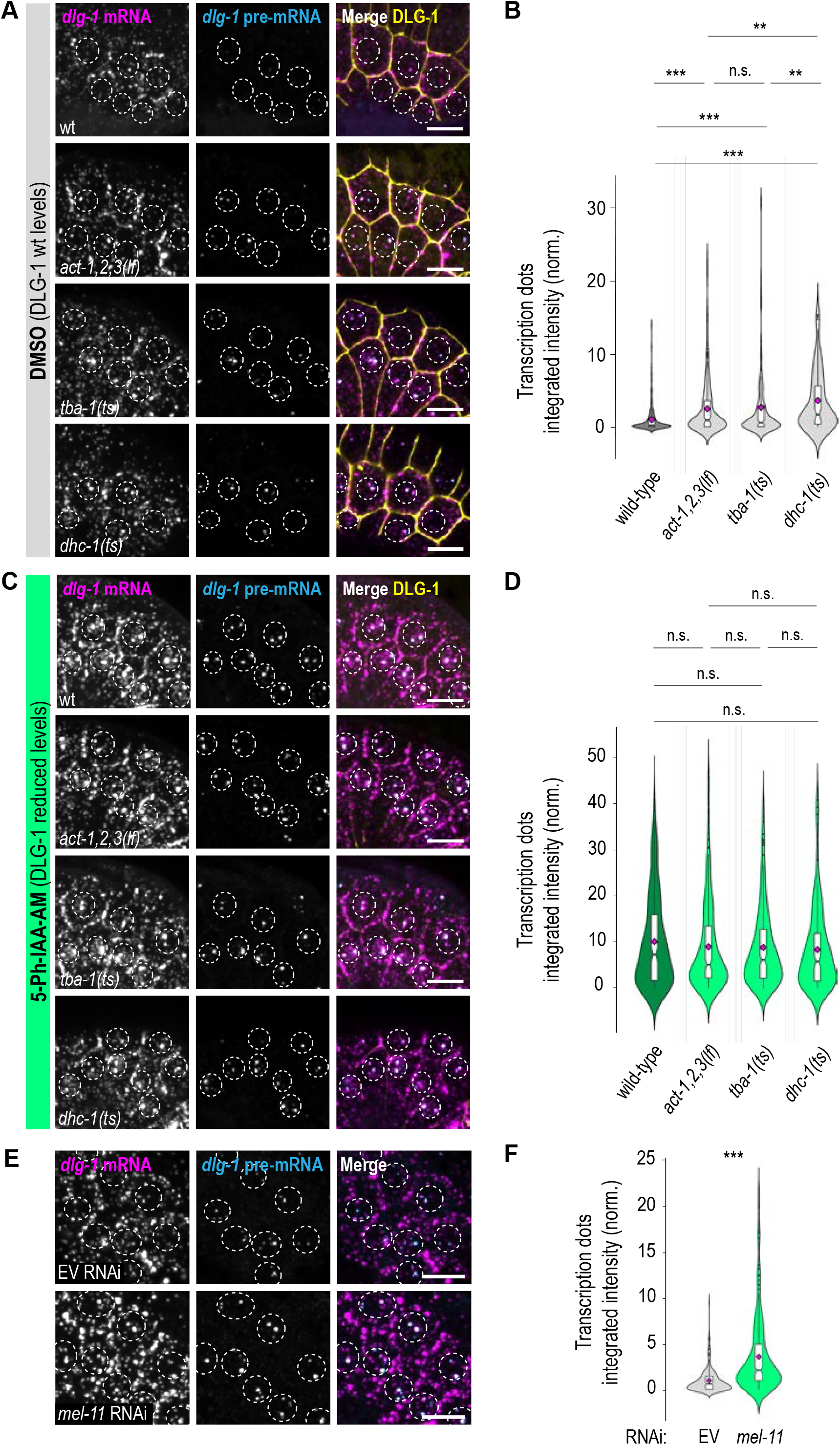
The cytoskeleton and mechanical forces are involved in transmitting the signal from the junction to the nucleus. **A.** Fluorescence images of epidermal cells of smFISH signal for *dlg-1* mRNA (magenta), *dlg-1* pre-mRNA (cyan), and merge with DLG-1::EGFP (yellow). All the strains shown in this panel contain endogenous *dlg-1* tagged with EGFP and mIAA7, but in different backgrounds and have been treated with DMSO (grey) for 30’. From top to bottom: wild type (wt); *act-1,2,3* hypomorphic mutant (*act-1,2,3(lf)*); *tba-1* thermo-sensitive mutant (*tba-(ts)*); *dhc-1* thermo-sensitive mutant (*dhc-(ts)*). Animals were grown on NGM plates with OP50 bacteria as food at 15°C, shifted for 2 hours at 24°C. Embryos were then collected, split into two: one half treated with DMSO (grey) for 30 minutes in tubes. Example nuclei of seam and ventral epidermal cells are circled with dashed lines. Scale bars: 5 µm. **B.** Violin plots with overlaid notched boxplots of distributions of integrated intensities of transcription dots (channel for *dlg-1* pre-mRNA) in the four strains shown in (A). n.s. = p > 0.05; ** = p < 0.01; *** = p < 0.001. For raw data and statistics, see Table S4. **C.** Fluorescence images as described in (A). In this case, embryos were treated with 5-Ph-IAA-AM (green) for 30’. Example nuclei of seam and ventral epidermal cells are circled with dashed lines. Scale bar: 5 µm. **D.** Violin plots with overlaid notched boxplots of distributions of integrated intensities of transcription dots (channel for *dlg-1* pre-mRNA) in the three strains shown in (C). n.s. = p > 0.05. For raw data and statistics, see Table S4. **E.** Fluorescence images of epidermal cells of smFISH signal for *dlg-1* mRNA (magenta) and *dlg-1* pre-mRNA (cyan), and merge with DLG-1::EGFP (yellow). The strain shown in this panel is the wild type (N2), on two RNAi treatments: upper panel: EV; lower panel: *mel-11*. Example nuclei of seam and ventral epidermal cells are circled with dashed lines. Scale bars: 5 µm. **F.** Violin plots with overlaid notched boxplots of distributions of integrated intensities of transcription dots (channel for *dlg-1* pre-mRNA) shown in (E). *** = p < 0.001. For raw data and statistics, see Table S4.

In a converse experiment, we induced actomyosin hypercontraction within developing embryonic epidermal cells using a mutation in the myosin-associated phosphatase orthologue *mel-11/PPP1R12A* (protein phosphatase 1 regulatory subunit 12A). Reduction of *mel-11* levels via RNAi induced *dlg-1* transcriptional upregulation (Fig.4E,F and S9A), indicating that J-QC is responsive to cytoskeletal disruptions, leading to transcriptional upregulation. Because *mel-11-*induced hypercontraction affects both the microtubule network and the actin cytoskeleton, we cannot determine which cytoskeletal network is primary^37,38^ In sum, these data suggest that the cytoskeleton is an integral component of the J-QC pathway controlling *dlg-1* transcription. The lack of an additive effect upon simultaneous disruption of DLG-1 and the cytoskeleton indicates that junctional and cytoskeletal integrity are monitored through the same genetic pathway.

### ZYG-12 mediates the J-QC response

We next considered how the non-nuclear compartment of the cytoskeleton communicates to the nucleus. The linker of nucleoskeleton and cytoskeleton (LINC) complex transduces signals between the cytoplasm and nucleus, ultimately remodeling gene expression during cell migration and development^39–43^. We therefore tested whether LINC components were required for the J-QC response.

LINC complexes consist of two interacting transmembrane proteins in the perinuclear space: a SUN (Sad-1-UNC-84) domain-containing protein of the inner nuclear membrane that connects to the nuclear lamina, and a KASH (Klarsicht, ANC-1, Syne homology) protein of the outer nuclear membrane that ultimately links the nucleus to the cytoskeleton. *C. elegans* possesses three SUN/KASH combinations that connect the cytoskeleton to the nucleus (Fig. 5A): SUN-1/ZYG-12 and UNC-84/UNC-83, which associate with microtubules through the motor protein dynein, and UNC-84/ANC-1, which binds actin^44,45^. These complexes have been studied in the context of nuclear migration and chromosome dynamics during germline and embryonic development in *C. elegans*^46,47^, but their roles in signal transduction and transcription remain unknown.

**Figure 5.**
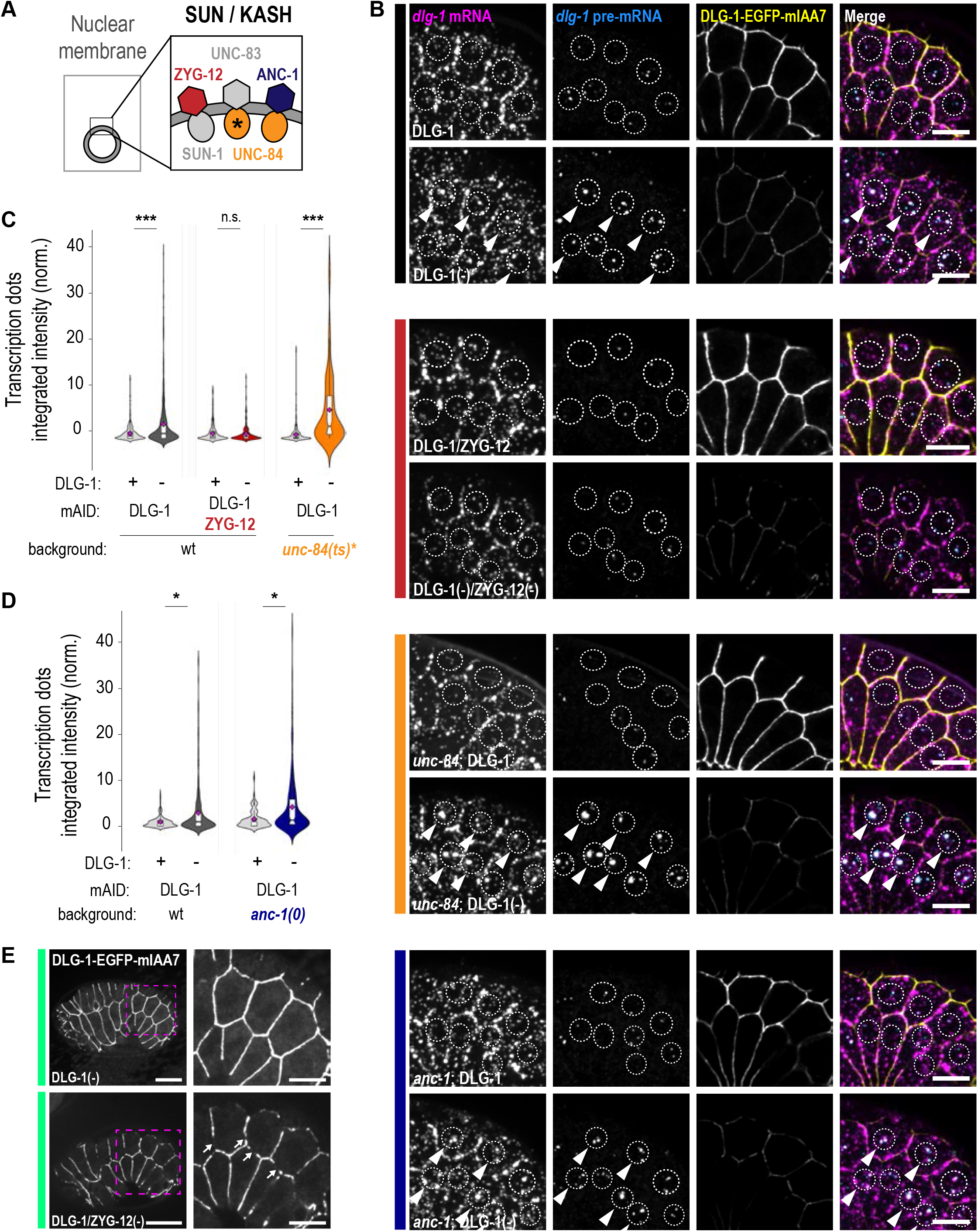
ZYG-12 is required to upregulate *dlg-1* in the J-QC mechanism. **A.** Simplified schematic representation of the three known LINC complexes in *C. elegans*. From left to right (KASH/SUN): ZYG-12/SUN-1, UNC-83/UNC-84*, and ANC-1/UNC-84. ZYG-12 (red), UNC-84* (orange with asterisk), and ANC-1 (dark blue) have been tested to cover the three respective complexes. **B.** Fluorescence images of epithelial cells of smFISH signal for *dlg-1* mRNA (magenta) and *dlg-1* pre-mRNA (cyan), DLG-1::EGFP::mIAA7 fluorescent signal (yellow), and merge. Embryos were treated with DMSO or 5-Ph-IAA-AM treatment for 30 minutes. From top to bottom: strain containing DLG-1 tagged with EGFP and mIAA7 (black vertical line on the side), treated with DMSO (DLG-1) (first raw) or with 5-Ph-IAA-AM (DLG-1(-)) (second raw); strain containing DLG-1 and ZYG-12, both tagged with green fluorophores and AID tags (red vertical line on the side), treated with DMSO (DLG-1/ZYG-12) (third raw) or with 5-Ph-IAA-AM (DLG-1(-)/ZYG-12(-)) (fourth raw); strain containing DLG-1 tagged with EGFP and mIAA7 in an *unc-84* mutant background constituted by an in-frame deletion (allele *e1174*) affecting specifically the interaction with UNC-83 and not ANC-1^93^ (orange vertical line on the side), treated with DMSO (*unc-84*;DLG-1) (fifth raw) or with 5-Ph-IAA-AM (*unc-84*;DLG-1(-)) (sixth raw); strain containing DLG-1 tagged with EGFP and mIAA7 in an *anc-1* mutant background (dark blue vertical line on the side), treated with DMSO (*anc-1*;DLG-1) (seventh raw) or with 5-Ph-IAA-AM (*anc-1*;DLG-1(-)) (eighth raw). Example nuclei are circled with dashed lines. Arrowheads point at example nuclei with large transcription dots. Scale bars: 5 µm. **C.** Violin plots with overlaid notched boxplots of distributions of integrated intensities of transcription dots for *dlg-1* pre-mRNAs in the different conditions. Data normalized to the control strain, DLG-1 tagged with mIAA7 and treated for 30 minutes with DMSO. n.s. = p > 0.05; *** = p < 0.001. For raw data and statistics, see Table S4. **D.** Violin plots with overlaid notched boxplots of distributions of integrated intensities of transcription dots for *dlg-1* pre-mRNAs in the different conditions. Data normalized to the control strain, DLG-1 tagged with mIAA7 and treated for 30 minutes with DMSO. * = p < 0.05. For raw data and statistics, see Table S4. **E.** Live fluorescence images of epidermal cells and zoom-ins (dotted magenta squares) for DLG-1::EGFP::mIAA7 signal upon 5-Ph-IAA-AM-mediated depletion of DLG-1 alone (DLG-1(-), upper panels) or both DLG-1 and ZYG-12 (DLG-1(-)/ZYG-12(-), lower panels). The images are the same as in Fig. 5B, but with higher contrast to focus on gaps at junctions (arrows). Scale bars: 10 μm (left panels) and 5 μm (right panels).

To test if SUN/KASH complexes were required for the J-QC, we paired DLG-1 degradation (the J-QC trigger) with loss of SUN/KASH components. Specifically, we disrupted the SUN-1/ZYG-12 and UNC-84/UNC-83 complexes, which connect the nucleus to microtubules, as well as the UNC-84/ANC-1 complex, which links the nucleus to actin^44,45^. To circumvent nonspecific effects by LINC component mutations, we used conditional (AID2 for *zyg-12*; thermo-sensitive *unc-84*) or loss-of-function (*anc-1*) alleles to test their function during morphogenesis. We found that loss of ZYG-12, but not UNC-84 or ANC-1, prevented the compensatory increase in *dlg-1* transcription observed upon DLG-1 depletion (Fig. 5B-D and S10A-E). Thus, ZYG-12 is required to induce the J-QC.

Our model suggests that loss of the J-QC will enhance phenotypes where the DAC has been compromised. We performed a double depletion of DLG-1 and ZYG-12 to disrupt the DAC and inhibit the J-QC response simultaneously. As expected, double inactivation produced worse junctional defects, including reduced DLG-1 protein and gapping along the DAC compared to DLG-1 inactivation with an active J-QC system (Fig. 5E). These results suggest a specific role for ZYG-12 in the J-QC response and underscore the importance of this system in responding when the DAC is compromised.

## Discussion

In this study, we identify two complementary processes that cooperate to ensure epithelial junctions remain intact. First, we show that subcellular localization of *dlg-1* mRNA promotes local DLG-1 protein accumulation at junctions; loss of mRNA localization leads to reduced DLG-1 and discontinuity within the DAC (gaps). Second, we uncover a novel quality control pathway, the junctional QC (J-QC), that triggers compensatory transcriptional upregulation of DAC components when the junction is compromised, and a signal is transduced to the nucleus by ZYG-12 (Fig. 6). Phenotypically, loss of the J-QC also leads to epithelial gaps, underscoring its importance. These findings suggest that epithelial junctions maintain homeostasis through the coordinated action of local protein production and adaptive transcriptional feedback. We suggest that spatially localized mRNAs enrich key structural proteins at sites of junction assembly, while the J-QC pathway adjusts transcriptional output in response to junctional integrity rather than sequence complementarity. These dual mechanisms ensure tissue integrity in the face of junctional perturbations.

**Figure 6.**
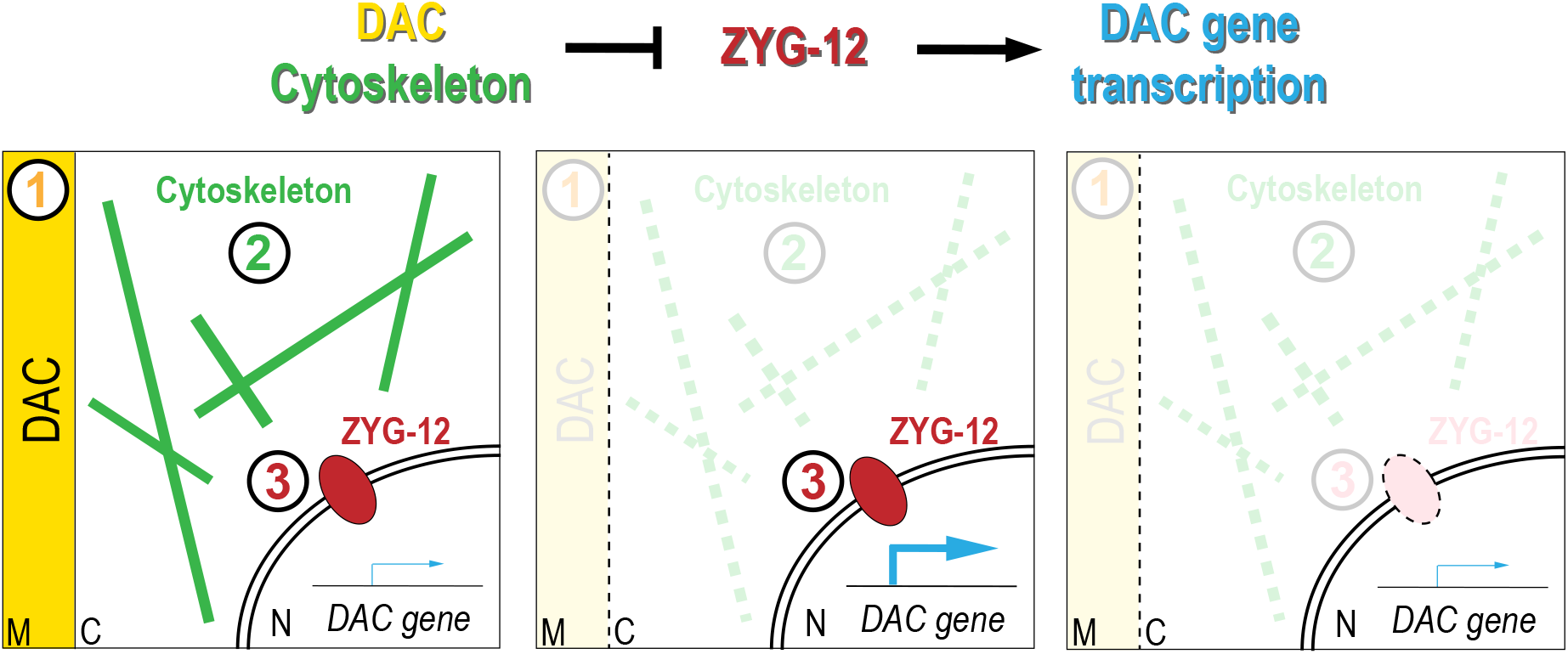
Model of the J-QC pathway. Proposed pathway in which the DAC components (yellow) and the cytoskeleton (green) negatively regulate ZYG-12 (red), which in turn positively regulates the expression of DAC genes (blue). Simplified schematic representation of the pathway within the cell. Left panel: under wild-type conditions, functional junctional DAC (yellow, #1), cytoskeleton (green, #2), and ZYG-12 (red, #3) maintain a basal level of transcription of junctional components (blue thin arrow). Middle panel: loss of DAC or cytoskeletal function triggers transcriptional upregulation of DAC genes (blue thick arrow). Right panel: transcriptional upregulation of DAC genes does not occur when ZYG-12 is lost in addition to the DAC.

### Conservation of DLG-1 orthologs and implications for disease

Our findings may be relevant to DLG-1 orthologues (DLG1) in other species and cell types. DLG-1 belongs to a highly conserved family of scaffolding proteins whose orthologs perform essential functions at cell-cell contacts across metazoans, including epithelial junctions^48^, neuronal synapses^49^, and immunological synapses in T-cells^50,51^. In several tumor types, total DLG1 protein is elevated, even if it is depleted from cell-cell contacts and redistributed to the cytoplasm. In cervical cancer, for example, DLG1 protein is both increased and preferentially cytoplasmic ^52^ and in in colorectal cancer, elevated cytoplasmic DLG1 protein correlates with reduced overall survival^53^. These patterns may reflect a J-QC–like response: cells sense the loss of junctional DLG1 and upregulate transcription in response, but when the additional protein fails to reach the junction, it drives further upregulation. High levels of cytoplasmic DLG1 correlate with poor prognosis in these cancers, suggesting this regulatory loop may be functionally important. Understanding the J-QC may offer a means to dismantle DLG1 mis-regulation in diseased states.

### mRNA localization as a component of junctional homeostasis

mRNA localization is widespread across species and cell types, with hundreds to thousands of localized transcripts identified in diverse biological systems^22,54,55^. In the best-understood paradigm, *cis*-regulatory “zip-code” elements, located in the 3’UTR, are recognized by RNA-binding proteins (RBPs) that direct the transcript to its destination^56,57^. RBP-bound transcripts are transported as part of larger, condensate-like ribonucleoprotein granules along microtubules, exemplified by kinesin-dependent transport of *oskar* mRNA^58^. This mechanism is particularly useful in situations where mRNA localization is temporally decoupled from translation, such as quiescent germ cells or highly polarized neurons. Classical studies established that maternally deposited mRNAs, such as *bicoid* and *oskar*, direct spatially restricted translation and protein localization after the onset of embryogenesis^59,60^, and this concept has since expanded across cell types and developmental stages^54,55,61–64^. Epithelia are another tissue rich in localized RNAs, but one where translation and mRNA localization are coupled temporally^22,65–68^. Here, we focus on epithelial *dlg-1*, which relies on a co-translational mechanism for mRNA localization rather than a UTR-based zip-code^22^. Currently, the best studied examples of translation-dependent mRNA localization are proteins that reside in membranous organelles such as the ER or mitochondrion, where translation kinetics and the size of the protein influence mRNA localization^69,70^. This pathway allows proteins to pass the membrane surrounding the organelles, but it is not binary, with a cohort of mRNA translated away from its destination. This observation raises the question of how important mRNA localization is for structures that lack a membrane. Our data with *dlg-1* show that mRNA localization has a large influence on integrity of the junction, by ensuring adequate levels of DLG-1 protein.

The functional consequence of localization differs between transcripts and species. In mouse melanoma cells, *Kif1c* mRNA localization is dispensable for total protein abundance but required for directed cell migration, reflecting a role in cellular polarity rather than protein levels^57^. Similarly, *C. elegans erm-1* mRNA localization is not required for protein abundance, but rather for intestinal morphogenesis^25^, implying that localization has roles beyond protein abundance. *dlg-1* transcripts represent the converse case: mRNA re-localization reduces total junctional DLG-1 protein, indicating that localization safeguards protein abundance at the site of junction assembly, but is not required for polarity *per se*. The DAC is incorporated into the junction only after the CCC and core polarity regulators are already established^19,21^. Therefore, junctional mRNA localization likely serves a reinforcing, structural role rather than the patterning function classically described to localized maternal transcripts. We suggest *dlg-1* mRNA remains translationally competent when directed to pores^25^, but that one or more events occur: i) the translated protein is not efficiently delivered to the junction; ii) specialized regulators selectively augment DLG-1 translation at the junction. Either of these hypotheses is consistent with the co-translational requirement for *dlg-1* mRNA localization^22^; iii) re-localized *dlg-1* transcripts or protein may undergo rapid degradation. Selective degradation has been described for other developmentally regulated transcripts through RBP/deadenylation- or miRNA-dependent mechanisms^71–73^. Thus, mRNA localization is a critical mode of regulation that ensures newly produced DLG-1 accumulates efficiently at the junction.

*dlg-1* orthologs (DLG1) are also subject to subcellular mRNA localization in vertebrate neurons^74,75^. This preferential subcellular localization does not depend on the *DLG1* 3’UTR^75^, suggesting a translation-dependent mode of localization analogous to what we discovered for *C. elegans dlg-1* in epithelia^22^. This observation suggests that translation-dependent mRNA localization, rather than RBP-mediated transport, may represent an evolutionarily conserved feature of *DLG1* transcripts. Because localization occurs while translation is already in progress, both the transcript and the protein synthesis machinery are brought to the specific subcellular locus. This mechanism allows fast local production of DLG1 protein and is particularly advantageous for structures that require rapid assembly or reinforcement, such as epithelial junctions and neuronal or immunological synapses. The work we presented here suggests that vertebrate neurons may also depend on mRNA localization to establish appropriate levels of DLG1 at the synapse. Our work provides a functional explanation for the conserved localization of *dlg-1* ortholog mRNAs, suggesting that they similarly support local DLG1 protein accumulation at synapses and other vertebrate cell-cell contacts.

### A structural QC mechanism for epithelial junctions

Our findings identify a novel QC system, the J-QC, that acts in epithelial cells to buffer junctional disruption. With the J-QC, cells couple the detection of structural perturbations of a defined subcellular entity to transcriptional responses to restore homeostasis. The J-QC is distinct from other QC mechanisms, which assess either organelle functionality or, alternatively, molecular fidelity. In the first instance, QC systems surveille membranous organelles such as mitochondria^8^, chloroplasts^12^ and the endoplasmic reticulum^13^ (ER). These systems commonly monitor the accumulation of damaged or misfolded proteins and restore organelle homeostasis through selective degradation mechanisms ^76–78^. In the second instance, systems monitor molecular processes such as DNA repair^79^, nonsense-mediated mRNA decay^80^ (NMD), transcriptional adaptation^33,34^, or the unfolded protein response^81^ (UPR). Transcriptional adaptation, for example, responds to NMD-mediated mRNA degradation by compensatory transcriptional induction of paralogous genes^35^. The importance of QC pathways is revealed by their wide ability to allow animals to survive compromised states, ranging from neurodegenerative disorders (*e.g*., Alzheimer’s disease) to disruptions in metabolism or protein-folding (*e.g*., type 2 diabetes and cystic fibrosis)^35,82–85^. Conceptually, the J-QC expands the scope of QC systems from monitoring proteins, RNAs, or membranous organelles to the maintenance of cellular architecture within epithelia.

A key feature of the J-QC is its remarkable specificity. Both trigger and response are restricted to the DAC: perturbation of DLG-1 or AJM-1 activates transcription of DAC genes, whereas disruption of the adjacent CCC or polarity regulators does not. This selectivity argues against a general stress response^86,87^ and instead suggests that epithelial cells distinguish between different junctional modules. Within the context of an organism, individual epithelial cell subtypes (*e.g*., the epidermis) may have evolved additional mechanisms to restrict transcriptional responses to specific genes, rather than activating a broad stress program. The precise transcriptional response sets the J-QC apart from other described forms of junctional surveillance, which act at the protein level but without altering gene expression: loss of tension across the CCC in mammalian models, for instance, triggers a conformational change in α-catenin and local recruitment of vinculin to reinforce the junction^88,89^. The J-QC instead couples a structural defect in one specific junctional module to a dedicated transcriptional program, revealing a module-selective layer of surveillance. Such specificity may allow epithelial cells to preserve tissue integrity while avoiding broad stress responses that could alter other aspects of development. More generally, our findings suggest that QC mechanisms can act not only on damaged proteins or organelles, but also on the composition of functional cellular structures whose precise organization is required during morphogenesis.

One possible explanation for the distinction between the DAC and the CCC is that the two junctional domains fulfill distinct roles. The cadherin-based CCC is established early during embryogenesis and provides the adhesive framework that maintains epithelial cohesion, whereas the Discs large-based DAC is built later, coinciding with the onset of morphogenesis and body elongation^19,21,32^. During this developmental transition, epithelial junctions experience continuous remodeling and variable mechanical forces^32,90^. A dedicated QC mechanism acting on the DAC could therefore represent a mechanism to reinforce junctions and maintain tissue integrity under variable mechanical load.

The J-QC operates independently of previously described RNA-mediated transcriptional compensatory pathways. The closest system is perhaps transcriptional adaptation in which mutated mRNAs induce transcriptional upregulation of paralogous genes^33,34^. However, J-QC is distinct from transcriptional adaptation since it does not depend on sequence similarity between the trigger and the responder, and it does not rely on nonsense-mediated decay components or premature translational termination. J-QC is also distinct from promoter-directed RNA activation (RNAa), which relies on transcriptional activation mediated by small RNA complementary to promoter regions^91,92^. Together, these observations indicate that the J-QC is not driven by RNA sequence or RNA metabolism but is a new type of surveillance system that monitors the state of the epithelial junction itself.

### The LINC component ZYG-12 mediates the transcriptional response in the J-QC pathway

The LINC complex was first genetically defined in *C. elegans* through the identification of SUN-and KASH-domain proteins required for nuclear anchorage and migration^93–95^. These studies established the LINC complex as a conserved functional unit that physically connects the cytoskeleton to the nucleus^39^. In vertebrate models, LINC complexes have been implicated in force transmission, particularly in nuclear positioning and mechanotransduction^87^. Actin-associated LINC complexes, such as nesprin-1/2 coupled to SUN2, participate in mechanotransduction pathways that regulate gene expression in response to changes in cellular tension and extracellular environment^96,97^. By contrast, microtubule- and dynein-associated LINC modules have primarily been linked to nuclear and chromosome positioning^98^, including the *C. elegans* KASH-domain protein ZYG-12^99–102^. However, whether distinct LINC modules can transmit specific information about cellular architecture to the nucleus to regulate transcription remains unclear. Our findings identify a previously unrecognized function for ZYG-12 in coupling epidermal junction defects to transcriptional upregulation of DAC components in the J-QC. Rather than representing a generic consequence of disrupting nuclear–cytoskeletal connections, this response requires a specific LINC module, suggesting that distinct LINC assemblies may have specialized roles in linking cellular architecture to nuclear responses.

ZYG-12 is a conserved member of the HOOK family of dynein adaptors. It combines a HOOK domain predicted to bind dynein light chain^103^, with a KASH domain that allows it to span the outer nuclear membrane and bind other components of the LINC complex. Together, these features enable ZYG-12 to couple microtubule-based transport through dynein to the nucleoplasm. Consistent with a role for dynein-dependent transport in this pathway, we showed that altering the function of dynein heavy chain (DHC-1) also affects the transcriptional response in the J-QC pathway, further supporting a functional connection between ZYG-12 and dynein-dependent transport processes. Interestingly, disruption of the actin cytoskeleton also activated the J-QC in addition to the microtubule- and dynein-associated LINC module. One possible explanation is that perturbation of the actin network indirectly alters microtubule organization. Indeed, actin and microtubules are extensively interconnected, and changes in actomyosin organization or contractility have been shown to remodel microtubule architecture^104^. Stil, we cannot completely exclude the possibility that both cytoskeletal systems may influence the J-QC directly.

Our analyses suggest a genetic pathway in which junctional DAC components and the cytoskeleton negatively regulate ZYG-12, which in turn positively regulates the expression of DAC genes (Fig. 6). Genetically, this model is supported by our finding that ZYG-12 is required for the transcriptional upregulation of DAC genes following disruption of either the DAC or the cytoskeleton, placing ZYG-12 downstream of these structural components and upstream of the transcriptional response. At the molecular level, our data are consistent with a model in which the integrity of the DAC and cytoskeleton is continuously monitored through a ZYG-12-dependent LINC module. Perturbation of this structural connection would then alter the signaling state of the pathway, leading to compensatory transcription of DAC genes. This model provides a unifying explanation for why diverse perturbations affecting junction integrity or cytoskeletal organization converge on the same transcriptional response, whereas disruption of adjacent junctional modules, such as the CCC, does not. We propose that the J-QC pathway functions to restore physiological levels of DAC components and thereby support epithelial biogenesis and embryonic elongation^105^.

### Methodology development

Our experiments relied on proven MS2/MCP methodology to re-localize *dlg-1* mRNA. Fusing MCP with nuclear pore components provides an alternative to RBP mutation or deletion strategies, which can induce pleiotropic effects unrelated to localization itself. Our approach is conceptually related to CRISPR-based tools such as CRISPR-TO^106^, CRISPR-TO technology uses dCas13 to reposition transcripts, which requires suitable guide sequences within the 3’UTR and assembling a multi-component complex that can interfere with gene expression independently of relocalization^106^. Our system is simpler as it requires only MS2 hairpins and a fused MCP protein, which can be targeted to any subcellular structure for which a suitable anchor protein exists. This system does not depend on a pre-existing localization signal or its cognate RBP, representing an advantage for transcripts like *dlg-1* that lack one. Importantly, re-localization was achieved directly in a live, developing organism rather than in cultured cells, making it possible to link mRNA localization directly to epithelial morphogenesis in the context of embryonic development rather than only to cellular phenotypes.

Quantitative analysis of transcription sites remains a challenge, particularly when comparing transcriptional activity across multiple genetic or experimental conditions. Although several tools have been developed for RNA spot detection and quantification^108,110^, DotQuant (https://github.com/angelo-angonezi/DotQuant) provides a streamlined workflow that integrates automated nuclear and three-dimensional spot segmentation and fluorescence measurements to enable rapid and reproducible measurement of transcription site intensity across large numbers of nuclei. While developed here for the quantitation of transcription sites in *C. elegans* embryos, the modular design of DotQuant can be adapted to other types of fluorescent puncta and imaging datasets from different systems. Furthermore, future incorporation of automated nuclear segmentation based on nuclear markers will extend its applicability by enabling fully automated selection and analysis of nuclei across different tissues and model systems.

## Methods

### Nematode culture

All animal strains were maintained as previously described (Brenner, 1974) at 20°C, unless stated otherwise. For experiments with temperature-sensitive strains, animals were grown at 15°C and shifted to the temperatures and times described in the corresponding figure legends and/or methods. RNAi experiments were performed as previously described except for the growing temperature which was set to 24°C, instead of 25°C, and starting from a synchronized population of molting L3-L4 animals left to grow until adulthood and lay eggs. Embryos were collected for further processing. For a comprehensive list of alleles, transgenic lines, and RNAi clones^107^, see Table S1.

### SUN/KASH alleles description

To perturb SUN-1/ZYG-12 function during morphogenesis, we used a strain carrying AID- and GFP-tagged ZYG-12 and induced degradation as previously described^99^, to bypass the early embryonic lethality of *sun-1* and *zyg-12* mutants. To perturb the UNC-84/UNC-83 complex, we used the temperature-sensitive *unc-84(n369ts)* allele (marked with an asterisk for clarity, *unc-84(ts)\**), which contains an in-frame deletion that specifically disrupts UNC-84 interaction with UNC-83 while preserving interaction with ANC-1^93^. To perturb the UNC-84/ANC-1 complex, we used the *anc-1(e1873*) mutant strain^95^. These perturbations were combined with the AID2 system for DLG-1::EGFP::mIAA7 degradation (to induce the J-QC) to analyze the J-QC response.

### Generation of transgenic lines

The transgenic strain where MCP was paired to NPP-9 was generated with the mosSCI technique^109^. Insertions were verified by genotyping, sequencing, and phenotypic readout. The transgenic sequences (promoter: *pes-10*, coding sequence: 3xFLAG fused to two MCP sequences, two mCherry sequences^23^, and the endogenous *npp-9* sequence (exons and introns); 3’UTR: *tbb-2*) were assembled with the NEBuilder® HiFi DNA Assembly Cloning Kit (New England BioLabs, cat#E5520), using pCFJ150 as a vector to insert the transgene into the *ttTi5605* locus on chromosome II (strain name: EG6699). The plasmid containing the transgene is called pCT3.82 and is available upon request. NEB® 10-beta Competent *E. coli* (High Efficiency) (NEB, cat#C3019) competent cells were used for transformation. Full plasmid sequencing from Microsynth was used to confirm the sequences were correct. Plasmids were purified with the NucleoBond Xtra Midi Plus (Macherey-Nagel, cat#740412.10) and single-copy integrated lines were generated and verified following the mosSCI protocol^109^.

The *dlg-1(Δc7aa)* embryos strain was generated with the CRISPR/Cas9 technique^111^ and consisted of a deletion of seven amino acids in the sequence linking SH3 and GuK domains of DLG-1 (aa:727-733). Two CRISPR RNAs (crRNA1: GAATGGGAACTTTCTGGAGA; crRNA2: CGGTCGGTACTTTTGACGAA; IDT), a single-stranded oligodeoxynucleotide as a repair oligo (ssODN, TGGAAAACCGAAGAGGAAGCCGCAGTCAGCTTTCTGTCAAAAGTACCGACCGGCTCAACGATCTTAATGA; IDT), Cas9 enzyme (Alt-R® S.p. Cas9 Nuclease V3, IDT, cat#1081058), and tracrRNA (IDT, cat#1072533) were used to generate the deletion strain on an already available allele where DLG-1 protein was tagged with mNG (LP598^28^). Genotyping was used to screen rollers, and further sequencing to verify correct deletion (Fw oligo: TGGATTGGAAAACCGAAGAG; Rv oligo: TGCTCAACTGCCTGGTAGG).

### smFISH

smFISH primary probes design was performed as previously described^22,112^. smFISH experiments were adapted from previously described protocols^22,112^. Adults from plates with a lot of laid embryos were removed by washing them off plates with distilled water (two times with 1 ml). The embryos and the leftover bacteria lawn were collected after adding 1 ml water and mechanically released by gentle rubbing with a gloved finger. Embryo suspensions were transferred to 1.5-ml microcentrifuge tubes and washed two times with distilled water by brief centrifugation (5 seconds, short pulse) to remove remaining worms, bacteria, and debris. Glass slides (Epredia™ Epoxy Diagnostic Slides, Fisher Scientific, cat#10393881) were coated with 1 µl poly-L-lysine and air-dried prior to use. Approximately 50 µl of embryo suspension was applied to each slide and allowed to settle for approximately 30 seconds. Excess liquid was removed, and embryos were immediately treated with fixative solution by adding 50 µl of freshly prepared 3.7% formaldehyde in PEM buffer for 5 minutes at room temperature in a humidity chamber. After removal of the fixative (though ensuring embryos did not dry), a small drop (2-3 µl) of the removed fixative was placed on a coverslip, which in turn was immediately applied to the slide. Slides were transferred onto a metal plate placed on dry ice and frozen for at least 30 minutes. Coverslips were removed by freeze cracking. Embryos were transferred in different coplin jars containing the following reagents: ice-cold methanol for 5 minutes, PBS for 1 minute at room temperature, 3.7% formaldehyde in PEM buffer for 5 minutes at room temperature in a second fixation step. Samples were briefly washed at room temperature in coplin jars as follows: 1 minute in PBS, 10 minutes in PBS-T, 20 minutes in PBS-T, and 5 minutes in PBS. Slides were dried but the well containing the embryos, which were with 100 µl hybridization solution for 1 hour at 37°C in a humidified chamber. Afterward, the blocking solution was removed and replaced with 100 µl hybridization solution containing smFISH probe(s) (2 µl per 100 µl hybridization solution for each FLAP duplex probe set, or 1 µl for Stellaris probes). Samples were incubated for at least 4 hours or overnight at 37°C in a dark, humidified chamber. Primary antibody against AJM-1 (MH27^113^, Developmental Studies Hybridoma Bank – DSHB) was added during this step when required. Following hybridization, samples were washed twice with wash buffer (WB) – 100 µl per wash by pipetting up and down on the well containing the sample. Where applicable, secondary antibodies (Goat anti-Mouse IgG (H+L) Highly Cross-Adsorbed Secondary Antibody, Alexa Fluor™ 488; Invitrogen, cat#A-11029) or nanobodies (GFP-Booster ATTO488, Chromotek, cat#gba488; RFP-Booster ATTO594, Chromotek, cat#rba594) diluted in WB were added and incubated for 1 hour at 37°C in the dark. Nuclei were stained with Hoechst (1:50 in WB) for 15 minutes at room temperature in a humidity chamber in the dark, followed by three washes in WB - 100 µl per wash by pipetting up and down on the well containing the sample. Samples were mounted in 12 µl of VECTASHIELD® Antifade Mounting Medium (Vector Laboratories; cat#H-1000-10) and sealed with nail polish prior to imaging and stored at 4°C in the dark.

Composition of buffers and solutions: Fixative solution (3.7% formaldehyde in 1xPEM buffer): formaldehyde solutions were prepared fresh from 37% stock. For on-slide fixation, 50 µl 37% formaldehyde was mixed with 100 µl 5xPEM buffer and 350 µl H₂O. For the second fixation in a coplin jar, 5 ml of 37% formaldehyde was mixed with 10 ml 5xPEM buffer and 35 ml H₂O. 5xPEM buffer (pH 6.8): 500 mM PIPES, 5 mM EGTA, 5 mM MgCl₂. PBS-T: PBS with 0.1% (v/v) of Tween-20. Wash buffer (WB): 10% (v/v) of formamide and 2xSSC in H₂O (usually prepared 10 ml and stored at 4°C). Hybridization solution: 10% (w/v) of dextran sulfate, 10% (v/v) of formamide, and 2xSSC in H₂O; always prepared fresh.

For *dlg-1*, *ajm-1*, *hrm-1*, *hmp-1*, *par-6*, and *pkc-3* mRNAs and *dlg-1* pre-mRNA we used probes previously described^22,23^. For the sequences of the newly made smFISH probes (*ajm-1* pre-mRNA), see Table S2.

### Microscopy

Images were acquired on embryos at the same stage (comma stage, at the onset of morphogenesis) and cell type (epidermal cells - dorsal-ventral and seam) throughout the manuscript for consistency.

For live imaging experiments, embryos were treated as previously described^22^. A widefield ZEISS Axio Imager M2 equipped with a ZEISS Colibri as LED light source, Hamamatsu ORCA flash 4.0 camera, and ZEN 2.6 software (blue edition) were used for capturing images for live imaging experiments. All images were processed in OMERO (https://www.openmicroscopy.org/omero/) and processed in the same manner within datasets. Figures were assembled in Adobe Illustrator (https://www.adobe.com/).

For imaging of fixed samples, in figures 1B and 2E a widefield microscope FEI “MORE” with total internal reflection fluorescence (TIRF), equipped with a Hamamatsu ORCA flash 4.0 cooled sCMOS camera, and a Live Acquisition 2.5 software were used for capturing images. smFISH pictures were then deconvolved with the Huygens software. In the rest of the figures, we used a Nikon Ti2 with perfect focus system (PFS), equipped with a CSU-W1 Yokogawa Spinning Disk Field Scanning Confocal System, a Hamamatsu ORCA-Fusion sCMOS camera, and a NIS elements with JOBS software to acquire images. All images were processed in OMERO (https://www.openmicroscopy.org/omero/), and figures were assembled in Adobe Illustrator (https://www.adobe.com/).

### Image analysis and quantitation

Quantitation of fluorescence signal from the junctions via live imaging: fluorescently labelled DLG-1 protein was quantified by analyzing the intensity (max) of five junctions connecting seam cells per embryo and removing the background signal (min). Fluorescent signal quantitation was performed in Fiji.

Quantitation of integrated intensities of transcription dots in smFISH experiments were performed via DotQuant (https://github.com/angelo-angonezi/DotQuant). We selected nuclei of interest following two approaches. In a cellpose-based approach, nuclei were segmented using Cellpose-SAM (https://doi.org/10.1101/2025.04.28.651001). Nuclei of interest were manually selected in Napari by identifying and recording the corresponding segmentation object IDs (https://napari.org/stable/) (Fig. S3A-B). Single-nucleus image crops were generated from the raw input images using the spatial coordinates defined by the selected segmentation masks. In a second, ROI-based approach regions of interest (ROIs) were manually defined around nuclei locations in Fiji using the ROI Manager tool (https://imagej.net/software/fiji/) (Fig. S3C-F). For each image, ROI coordinates were exported as a .csv file describing a three-dimensional bounding box specified by five parameters: center coordinates (X, Y, Z), width, and height (Fig. S3G). Single-nucleus image crops were generated using a custom Python script that integrated the raw image data with the corresponding ROI coordinate files. For each ROI, a fixed axial depth of 10 Z-slices (±5 slices from the ROI Z-center) was used (Fig. S3H). Cropped nuclei were post-processed using a Difference of Gaussians (DoG) filter to enhance small, high-intensity puncta. FISH signals were segmented by intensity thresholding of the DoG-filtered images (a different set of thresholds was applied to pre-mRNA and mRNA per dataset due to varying intensities between channels). Individual dot volumes were calculated from the voxel-based binary masks by converting each segmented object into a surface mesh and computing its volume using the Python trimesh library (https://pypi.org/project/trimesh/). In nuclei containing more than two detected dots, only the two largest objects (by volume) were retained for downstream analysis. To maintain a consistent dot count per nucleus, one or two empty dots (volume = 0.0, intensity = 0.0) were added to nuclei containing fewer than two detected dots. For intensity quantification, the segmentation masks obtained in the previous step were applied to the original images to extract the pixels corresponding to each individual dot. The mean intensity was then calculated for each dot. Background intensities used for normalization were calculated as the mean intensity within each single-nucleus crop, excluding pixels belonging to the dot masks. Integrated intensity was defined as the product of each dot rescaled mean intensity and the corresponding mesh volume.

### Statistical analyses and data visualization

Data analysis and visualization were performed in R version 4.5.1 (https://www.R-project.org/) using the packages ggplot2 (https://ggplot2.tidyverse.org) and dplyr (https://dplyr.tidyverse.org). The Shapiro–Wilk test was used to check normality of the data and verified with normality diagnostic plots for such a test. Homogeneity of variances was evaluated using Levene’s test. For normally distributed data with equal variances, a one-way ANOVA was performed. For normally distributed data with unequal variances a pair-wise *t*-tests with Holm correction for multiple comparisons was used (Fig. 1D,G, 2B,E). For non-normally distributed data, group differences were assessed using the Kruskal-Wallis test. When the Kruskal-Wallis test was significant, it was followed by pairwise Wilcoxon rank-sum tests with continuity correction (P value adjustment Benjamini-Hochberg for multiple testing correction) to test pairs (Fig. 2A,G, 3C,E,F, 4B,D,F, 5C,D). Bar plots were used for normally distributed data: the bar represents the median, whiskers the standard deviation (SD), diamonds the mean, and black dots individual data points. Violin plots with overlaid notched box plots were used for non-normally distributed data: median (central thick line), interquartile range (IQR, box), and approximate 95% confidence interval (whiskers). Notches represent the confidence interval (CI) around the median. Magenta diamonds indicate the mean, and dots represent outliers. Statistical significance is indicated above bars or notched boxes as specified in each figure legend. For raw data, sample size (n), mean, median, and SD from each experiment, see Table S3 (live samples) and Table S4 (fixed samples).

### mRNA re-localization

The *dlg-1* mRNA re-localization experiments were performed by adapting the MS2-MCP technique. Specifically, *dlg-1* mRNA with or without MS2 hairpins in its 3’UTR was paired to MCP fused or not to the nuclear pore protein NPP-9/NUP358. The four transgenic combinations were all in an NMD-deficient background (*smg-2/UPF1,* allele *e2008)* to prevent degradation of *dlg-1* mRNA containing MS2 hairpins^23^. NPP-9 is on the cytoplasmic surface of the nuclear pore, and *dlg-1* mRNAs containing MS2 hairpins were trapped in this subcellular location by the MCP::NPP-9 protein fusion upon exiting the nucleus. In this way, *dlg-1* mRNA could not reach its normal location at the apical junction. Experiments on the four strains were performed at 24°C to allow basal expression of MCP from the heat-shock promoter as previously described^23^.

### Protein degradation experiments with AID2 system

The DLG-1 degradation experiments were performed following the AID2 system that takes advantage of a modified TIR1 enzyme from *Arabidopsis thaliana* (AtTIR1(F79G)) and the eggshell-permeable 5-phenyl-indole-3-acetic acid acetoxymethyl ester (5-Ph-IAA-AM), a modified synthetic analog of the auxin hormone^30^. Laid embryos from the *C. elegans* strain containing the TIR1 enzyme and mIAA7-EGFP-tagged DLG-1^31^ were collected in an Eppendorf tube and resuspended in M9. Some embryos from this sample were used for live imaging and fluorescence quantitation analyses. The remaining sample was then split in half into two separate Eppendorf tubes: DMSO (Dimethyl sulfoxide, Sigma Aldrich, cat#D4540) was added to the control sample, and 5-Ph-IAA-AM (Tocris Bioscience™ 5-Ph-IAA-AM, Fischer Scientific, cat#18786115) (resuspended in DMSO) was added to the treated sample for a final concentration of 50 µM. The samples were placed on a rotating wheel and one third of embryos from both samples (DMSO and 5-Ph-IAA-AM) were collected at 15, 30, and 45 minutes for the time course experiment. These pools of embryos were then split and used for both live imaging for fluorescence quantitation analyses, and to prepare slides to be processed for smFISH experiments. In the other AID2 experiments, including the double DLG-1/ZYG-12 degradation, embryos were collected after a 30-minute treatment.

### Real-time quantitative PCR (RT-qPCR)

Total RNA was isolated from pelleted embryos using the RNeasy Plus Mini Kit (Qiagen, cat#74134) following the manufacturer’s protocol. Residual genomic DNA was removed by treatment with DNase I (Thermo Fisher Scientific, cat#18060-015). RNA concentration and purity were assessed spectrophotometrically. First-strand cDNA synthesis was performed using the SuperScript™ IV First-Strand Synthesis System (Thermo Fisher Scientific, cat#18091050) with random hexamer primers, using DNase-treated RNA as input. cDNA samples were diluted prior to quantitative PCR analysis. Real-time quantitative PCRs were performed using PowerUp™ SYBR™ Green Master Mix (Applied Biosystems, cat#A25742). Reactions were performed using 5 ng of cDNA per reaction and gene-specific primers at a final concentration of 500 nM in a final reaction volume of 10 µL. Amplification was performed on a real-time PCR system under standard cycling conditions, followed by melt-curve analysis to confirm amplification specificity. No-template controls were included in all experiments. Relative gene expression levels were determined using the comparative Ct (ΔΔCt) method. Ct values for target genes were normalized to those of a reference housekeeping gene (*ama-1*) to obtain ΔCt values. ΔΔCt values were calculated relative to the control condition, and fold changes in gene expression are shown in Fig. S7E. For reagents and raw data from the RT-qPCR experiments, see Table S5.

## Glossary

AID: auxin-inducible degron
CCC: cadherin-catenin complex
DAC: DLG-1-AJM-1 complex
MCP: MS2 coat protein
NMD: nonsense-mediated mRNA decay
NPP: nuclear pore protein
QC: quality control
J-QC: junctional quality control
RT-qPCR: real-time quantitative PCR
smFISH: single molecule fluorescent *in situ* hybridization

## Acknowledgments

We thank the Boxem and Gassmann groups for sharing respectively their DLG-1 and ZYG-12 strains paired to fluorophores and AID tag, the Imaging Core Facility of the Biozentrum for technical support, current and previous lab members of the Mango group and the Basel RNA Club for scientific discussions, and WormBase. Special thanks to Miriam Edmunds for technical support during data analysis. Some strains were provided by the CGC, which is funded by the NIH Office of Research Infrastructure Programs (P40 OD010440).

The study has been partially funded by the Forschungsfonds (Excellent Junior Researcher) of the University of Basel (U.570.1040) and the Spark grant of the Swiss National Science Foundation (CRSK-3_228899) held by CT and the SNSF grant (SNF 310030_197713) held by SEM.

## Supplementary figure legends

**Figure S1.**
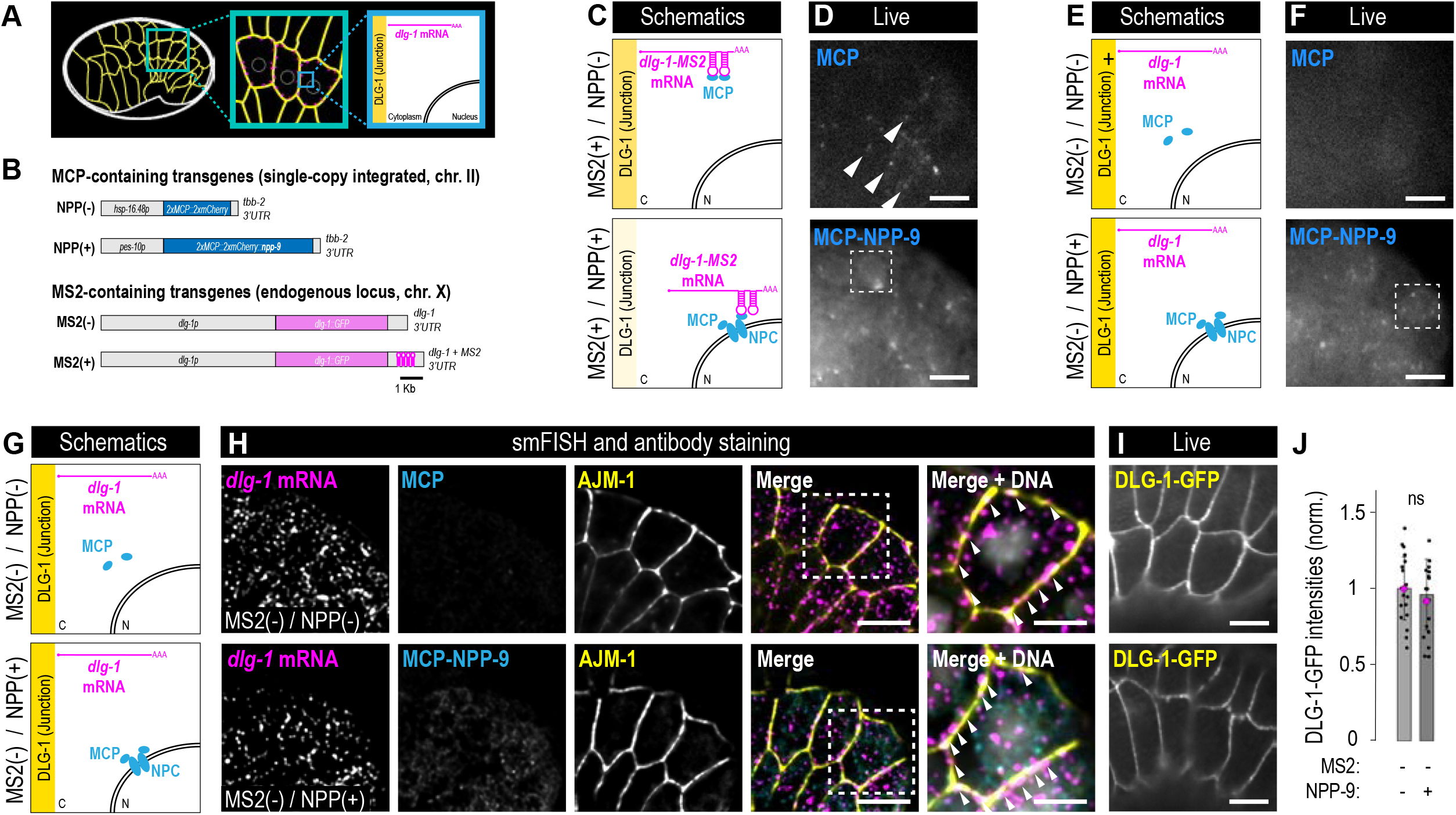
An adapted MS2-MCP method allows re-localization of MS2-containing transcripts. **A.** Left panel: schematic representation of a *C. elegans* embryo at the comma stage showing in yellow the junctions surrounding developing epidermal cells; blue square: portion of the embryo zoomed in in the middle panel. Middle panel: schematic representation of a zoom-in of a portion of *C. elegans* embryo in the left panel showing in yellow the junctions surrounding developing epidermal cells; dashed circles: nuclei; blue square: portion of the embryo zoomed-in in the right panel. Right panel: schematic representation of a portion of a seam cell from the blue square of the middle panel. Yellow: junctions (where DLG-1 localizes); magenta: *dlg-1* mRNA localizing at the proximity of the junction; nuclear membrane is also shown. **B.** Schematic representation of transgenes used in the re-localization experiments. MCP-containing transgenes without (NPP(-)) or with (NPP(+)) NPP-9 (blue) are expressed under the promoter of the early embryonic gene *pes-10*, and with the 3’UTR of the *tbb-2* gene. The transgene are single-copy integrations (mosSCI) on chromosome II. Endogenous *dlg-1* transgenic lines are CRISPR-derived. Their CDS are both tagged with in-frame GFP (magenta). One possessed endogenous 3’UTR (MS2(-)) and the other 24xMS2 hairpins (MS2(+)). Scale bar: 1 Kb. **C.** Schematic representation as in Fig. 1A. **D.** Live fluorescence images of a portion of *C. elegans* embryos at the comma stage, focusing on epidermal cells. Fluorescence signal in upper image: 2xMCP::2xmCherry bound to *dlg-1* containing MS2 hairpins in its 3’UTR; arrowheads point at dots representing examples of localized *dlg-1* transcripts at the junctions. Lower image: 2xMCP::2xmCherry::NPP-9 localized perinuclearly; dashed square: example of a nucleus surrounded by transgenic and fluorescently tagged NPP-9. Scale bars: 5 µm. **E.** Schematic representation of a portion of a seam cell as described in (Fig. 1A). Additionally, in both panels: endogenous *dlg-1* mRNA without MS2 hairpins (MS2(-), magenta). In cyan, upper panel: MCP alone (NPP(-)) not bound to junctional *dlg-1* mRNA and dispersed in the cytoplasm; lower panel: MCP fused to NPP-9 (NPP(+)) and not bound to junctional *dlg-1* mRNA and localizing at the NPC. **F.** Live fluorescence images of a portion of *C. elegans* embryos at the comma stage, focusing on epidermal cells. Fluorescence signal in upper image: 2xMCP::2xmCherry diffuse in the cell as *dlg-1* transcripts do not contain MS2 hairpins in the 3’UTR. Lower image: 2xMCP::2xmCherry::NPP-9 localized perinuclearly; dashed square: example of a nucleus surrounded by transgenic and fluorescently tagged NPP-9. Scale bars: 5 µm. **G.** Schematic representation as in (E). **H.** Fluorescence images of epidermal cells of smFISH signal for *dlg-1* mRNA (magenta), residual fluorescence signal of MCP (absent) and MCP::NPP-9 (perinuclear localization) (cyan), and antibody staining of AJM-1 (yellow). Merged images and zoom-ins of one seam cell (dotted square) are shown in the last two panels. Arrowheads point at transcripts (magenta) localized at the junction (yellow) in both conditions. Scale bars: 5 µm (first four panel) and 2.5 µm (last panel). **I.** Live fluorescence images of a portion of *C. elegans* embryos at the comma stage, focusing on epidermal cells. Fluorescence signal in upper image: DLG-1::GFP in MS2(-); NPP(-) background; lower image: DLG-1::GFP in MS2(-); NPP(+) background. Scale bar: 5 µm. **J.** Bar plot with error bars normalized to the control MS2(-); NPP(-) (light grey) and showing no reduction in DLG-1::GFP fluorescence intensities in MS2(-); NPP(+) (dark grey). Each dot represents the maximum fluorescent intensity of a junction shared between two seam cells (three junctions from the posterior-most seam cells per embryo) marked with endogenous DLG-1::GFP. n.s. = p > 0.05; *** = p < 0.001. For raw data and statistics, see Table S3.

**Figure S2.**
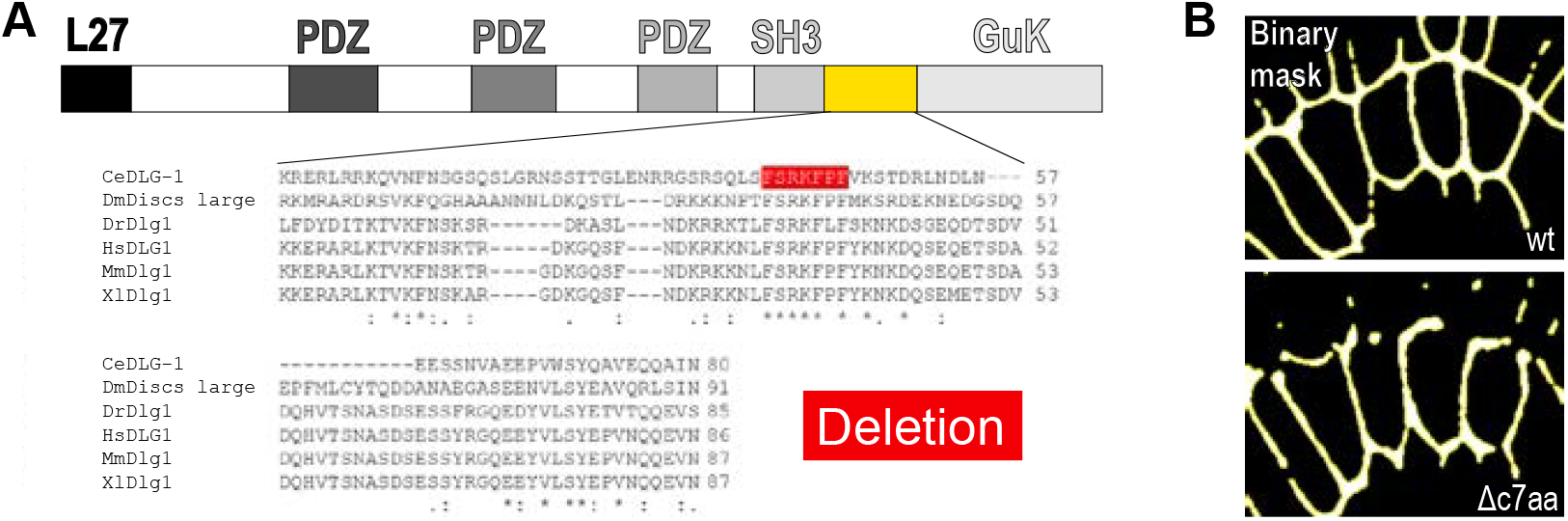
Deletion of seven amino acids between the SH3 and GuK domains of DLG-1 causes gap formation along the DAC. **A.** Schematic representation of DLG-1 domains (grey) with a focus on the sequence between SH3 and GuK (yellow). Sequence alignment (ClustalOmega) of DLG-1 Hook sequences from different species. From top to bottom: *C. elegans*, *D. melanogaster*, *Danio rerio*, *Homo sapiens*, *Mus musculus*, *Xenopus laevis*. Highlighted in red, the seven amino acids in a conserved region that have been deleted in the *dlg-1(Δc7aa)* strain. **B.** Binary masks in wild type (wt, upper image) and *dlg-1(Δc7aa)* embryos (Δc7aa, lower image) highlighting the presence of multiple gaps along the DAC of *dlg-1(Δc7aa)*.

**Figure S3.**
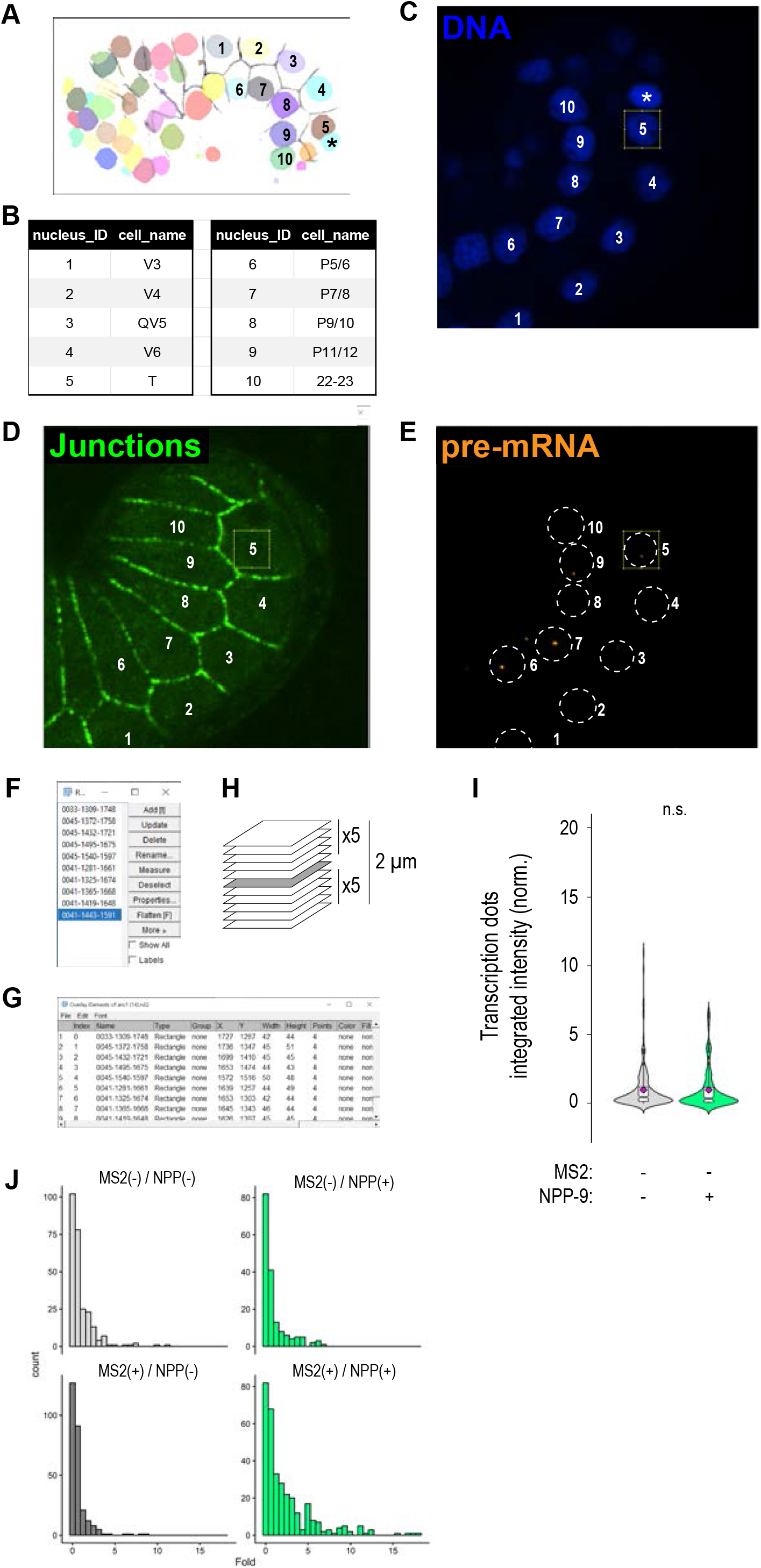
Integrated intensity analyses of *dlg-1* transcription sites via DotQuant reveal increased transcriptional levels upon mRNA re-localization. **A.** Segmentation of nuclei of a *C. elegans* embryo at the comma stage from a single stack and focusing on epidermal cells. Each identified nucleus is marked with a different color. In faded grey, junctions. The numbers indicate the nuclei used for the integrated intensity analyses of transcription sites throughout this study. X represents a reference nucleus used for cell identification. **B.** List of names of the cells whose nuclei are numbered in (A). **C,D,E.** DNA staining (Hoechst, blue, (C)), junctional staining (AJM-1, green, (D)), and transcription sites (intron smFISH probes for *dlg-1*, orange, (E)) showing the same nuclei/cells shown in (A) and listed in (B). A square shows a region of interest (ROI) to select the nuclei to analyze. **F.** List of the 10 ROIs selected on an image in Fiji. **G.** List of coordinates for each selected ROI. **H.** Schematics of Z stacks chosen for the analysis of transcription sites: the middle plane of a nucleus is chosen as ROI and in addition to it the 5 planes above and below are used for the quantitation of integrated intensity of the identified dots. **I.** Violin plots with overlaid notched boxplots of distributions of integrated intensities of transcription dots. n.s. > 0.05. For raw data and statistics, see Table S4. **J.** Histograms of distributions of fold values across strains from (I) and Fig. 2A. Fold values are on the x axis and binned into 30 intervals, displayed in separate panels for each strain. Bin counts are shown on the y axis, allowing comparison of distribution shapes within each condition.

**Figure S4.**
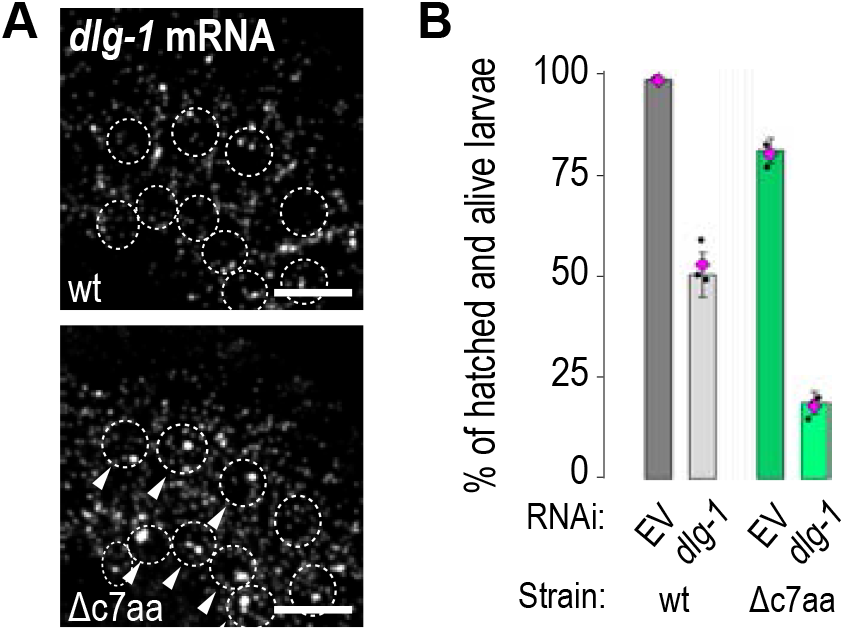
Reduced *dlg-1* mRNA levels exacerbate *dlg-1(Δc7aa)* embryonic lethality. **A.** Fluorescence smFISH images for *dlg-1* mRNA of epidermal cells of wild type and *dlg-1(Δc7aa)* mutant. Arrowheads point at nuclei (dotted circles) with larger transcriptional foci in *dlg-1(Δc7aa)* compared to the wild-type control. Scale bars: 5 µm. **B.** Bar plot with error bars showing the changes in embryonic viability in the different tested genetic conditions. Each dot represents the percentage of hatched and alive larvae per biological replicate (x3) in the different conditions. For raw data and statistics, see Table S3.

**Figure S5.**
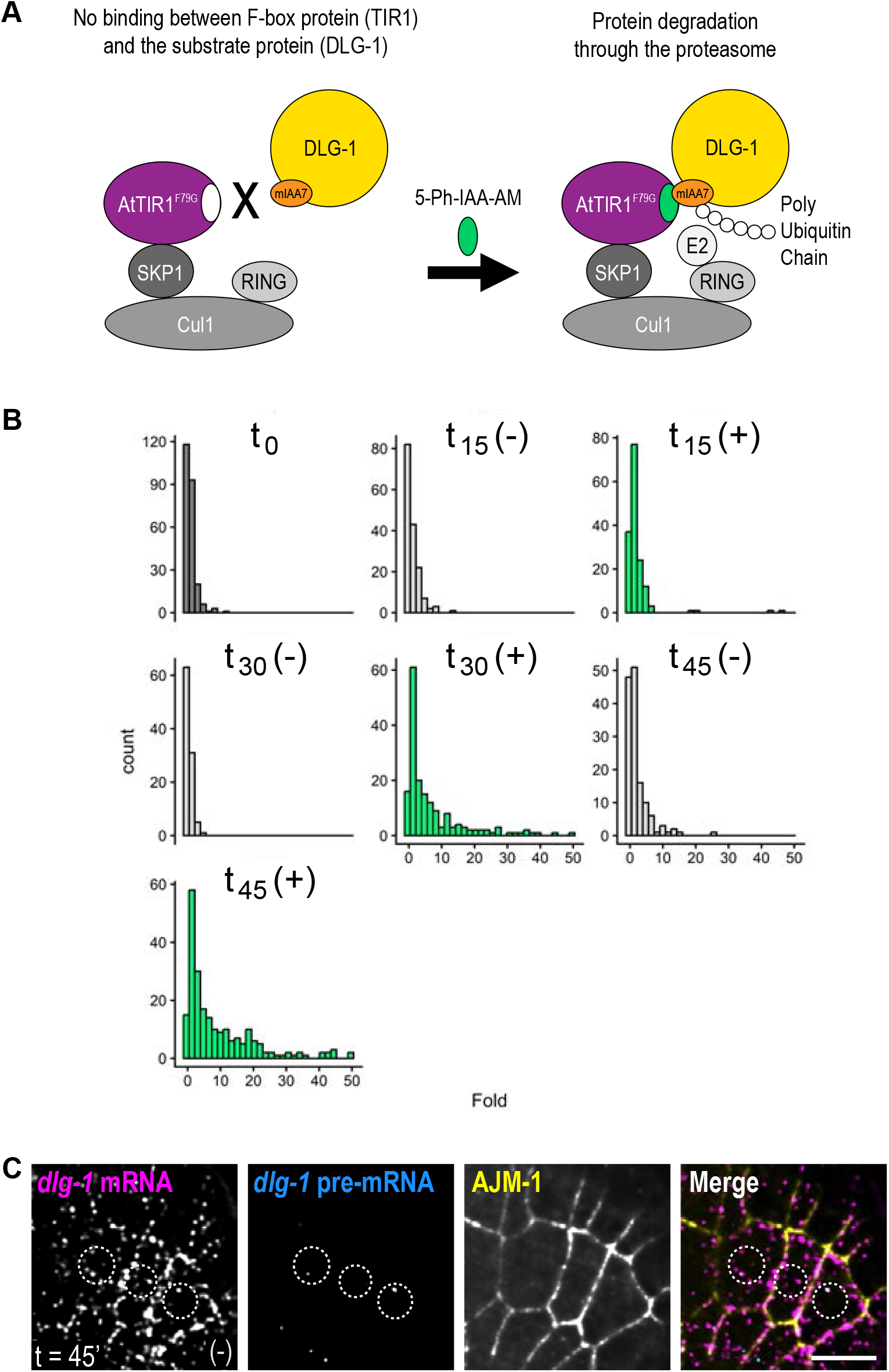
DMSO treatment does not cause *dlg-1* transcriptional upregulation after 45 minutes. **A.** Schematic representation of the AID2 system. The E3 complex (Cul1, SKP1, RING (shades of grey), and TIR1 (purple, mutant F79G from *Arabidopsis thaliana*)) cannot bind mIAA7-tagged DLG-1 (left). Upon addition (arrow) of 5-Ph-IAA-AM (green), AtTIR1(F79G) can bind mIAA7-tagged DLG-1 and allows its poly-ubiquitination by the E2 and sent for proteasome degradation (right). **B.** Histograms of distributions of fold values across conditions during the AID2 time course experiments. Fold values are on the x axis and binned into 30 intervals, displayed in separate panels for each strain. Bin counts are shown on the y axis, allowing comparison of distribution shapes within each condition. **C.** Fluorescence images of epidermal cells of smFISH signal for *dlg-1* mRNA (magenta), smFISH signal for *dlg-1* pre-mRNA (cyan), and antibody staining of AJM-1 (yellow) and merge. Example nuclei are circled with dashed lines. Scale bar: 5 µm.

**Figure S6.**
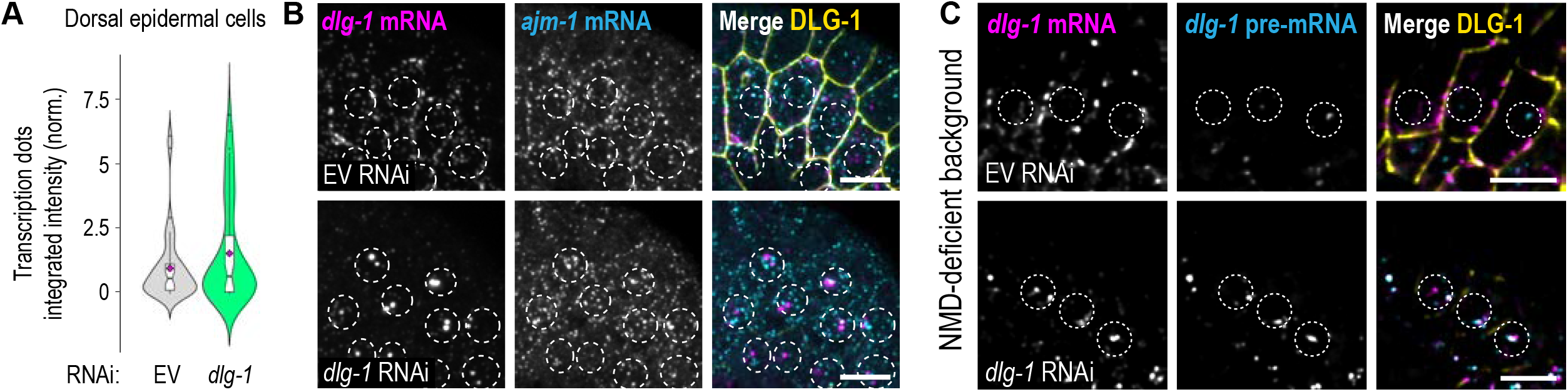
Reduction of DAC factors induces a transcriptional increase of DAC components and is independent from NMD. **A**. Violin plots with overlaid notched boxplots of distributions of integrated intensities of transcription dots for *ajm-1* pre-mRNAs from dorsal epithelial cells. Data normalized to the EV RNAi control. For raw data and statistics, see Table S4. **B.** Fluorescence images of seam cells. From left to right, the panels show smFISH signal for *dlg-1* mRNA (magenta), *ajm-1* mRNA (cyan), and merge with DLG-1::EGFP (yellow). Upper panel: EV RNAi. Lower panels: *dlg-1* RNAi. Example nuclei of seam and ventral epidermal cells are circled with dashed lines. Scale bar: 5 µm. **C.** Fluorescence images of a portion of fixed *C. elegans* embryos with an NMD-deficient background (*smg-2(e2008)*) at the comma stage, focusing on seam cells. From left to right, the panels show smFISH signal for *dlg-1* mRNA (magenta), *dlg-1* pre-mRNA (cyan), and merge with DLG-1::GFP (yellow). Upper panel: EV RNAi. Lower panels: *dlg-1* RNAi. Example nuclei of seam cells are circled with dashed lines. Scale bar: 5 µm.

**Figure S7.**
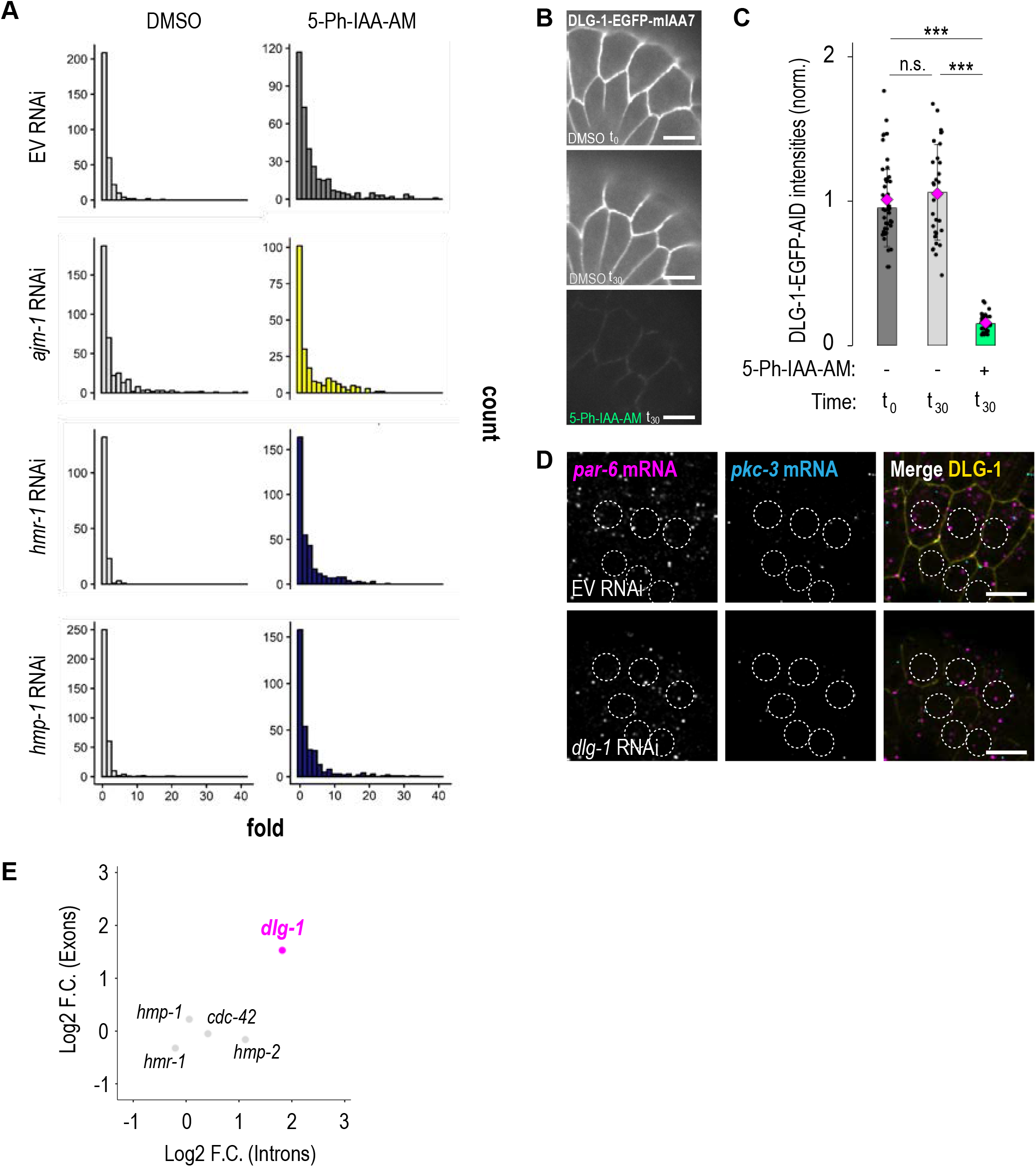
CCC and polarity regulators are not influenced by the J-QC. **A.** Live fluorescence images of a portion of *C. elegans* embryos at the comma stage, focusing on epidermal cells. Fluorescence signal of DLG-1::EGFP::mIAA7 strain. Upper image: treatment with DMSO at t0 (baseline control); middle image: treatment with DMSO at t30. Lower image: treatment with 5-Ph-IAA-AM at t30. Scale bar: 5 µm. **B.** Bar plot with error bars normalized to the reference t0 (dark grey) and showing the changes in DLG-1::EGFP fluorescence intensities using the AID2 system after 30 minutes (light grey: DMSO treatment; green: 5-Ph-IAA-AM treatment). Each dot represents the maximum fluorescent intensity of a junction shared between two seam cells (three junctions from the posterior-most seam cells per embryo) marked with endogenous DLG-1::EGFP. n.s. = p > 0.05; ***: p < 0.001. For raw data and statistics, see Table S3. **C.** Histograms of distributions of fold values across conditions. Fold values are on the x axis and binned into 30 intervals, displayed in separate panels for each strain. Bin counts are shown on the y axis, allowing comparison of distribution shapes within each condition. **D.** Fluorescence images of epidermal cells of smFISH signal for *par-6* mRNA (magenta), *pkc-3* mRNA (cyan), and merges with DLG-1::EGFP (yellow). Upper panel: EV RNAi. Lower panels: *dlg-1* RNAi. Example nuclei of seam and ventral epidermal cells are circled with dashed lines. Scale bars: 5 µm. **E.** Scatter plot of intronic (x axis) and exonic (y axis) reads of data derived from RT-qPCR (log₂ fold change, *ajm-1* mutant versus wild type). Each point represents a gene tested and shows the difference in expression levels between conditions, with *dlg-1* highlighted in magenta and all the other genes shown in grey.

**Figure S8.**
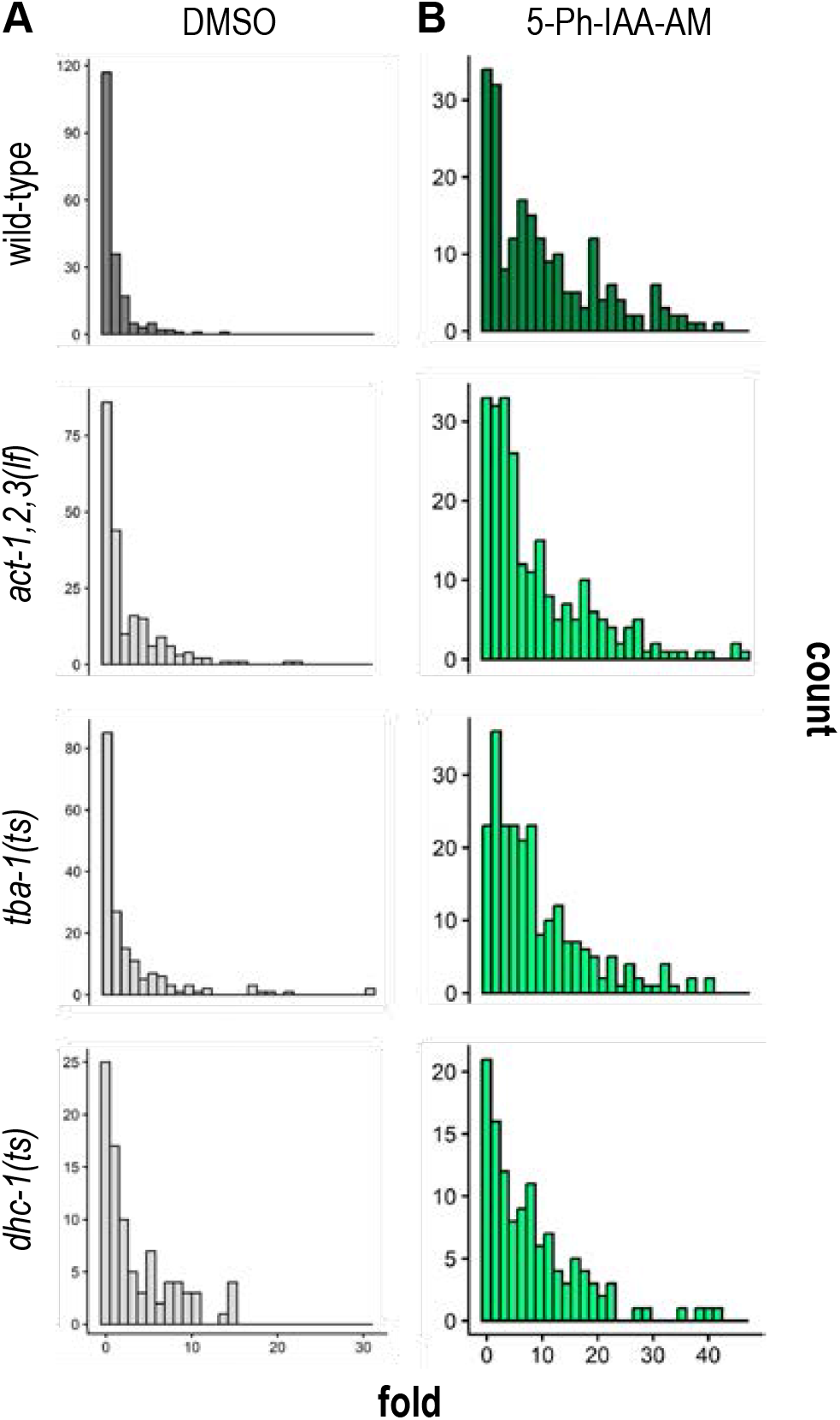
Different components of the cytoskeleton are involved in the J-QC mechanism. **A,B.** Histograms (grey data from Fig. 4B and green data from Fig. 4D) of distributions of fold values across strains and conditions. Fold values are on the x axis and binned into 30 intervals, displayed in separate panels for each strain. Bin counts are shown on the y axis, allowing comparison of distribution shapes within each condition derived from Fig. 4B.

**Figure S9.**
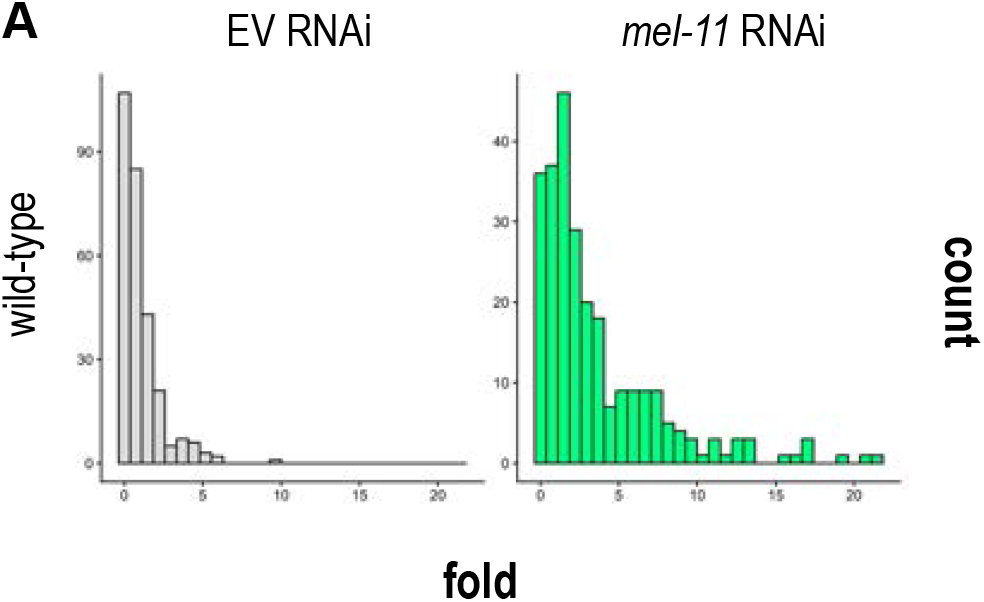
Mechanical forces are involved in the J-QC mechanism. **A.** Histograms of distributions of fold values across conditions from Fig. 4F. Fold values are on the x axis and binned into 30 intervals, displayed in separate panels for each strain. Bin counts are shown on the y axis, allowing comparison of distribution shapes within each condition: EV RNAi (grey) and *mel-11* RNAi (green).

**Figure S10.**
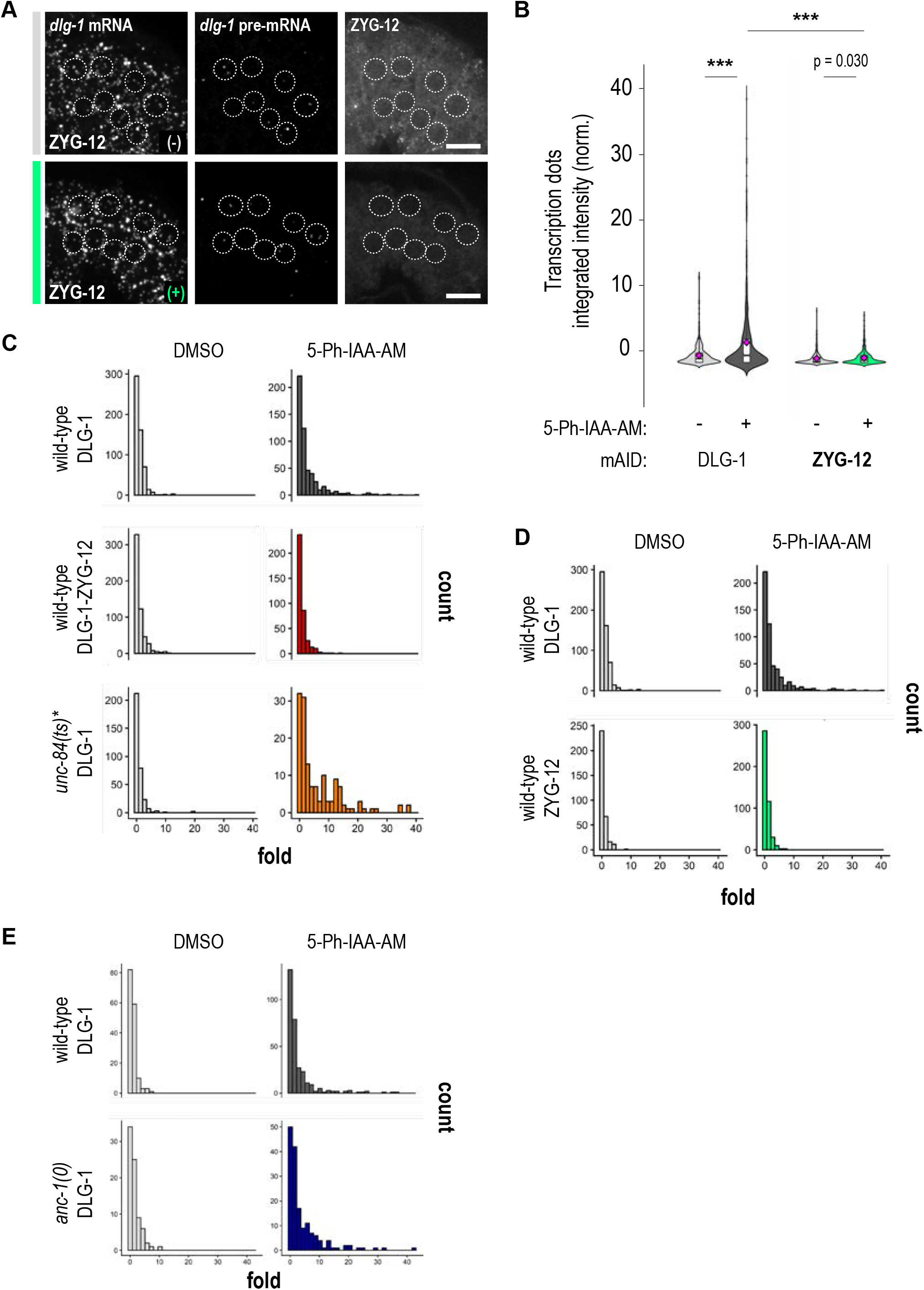
Removal of ZYG-12 does not impact *dlg-1* transcriptional regulation but inhibits the J-QC upon DLG-1 loss. **A.** Fluorescence images of seam and ventral epithelial cells. Upper panels: DMSO (grey). Lower panels: 5-Ph-IAA-AM (green). In both instances: treatment for 30 minutes. From left to right: smFISH signal for *dlg-1* mRNA, smFISH signal for *dlg-1* pre-mRNA, and ZYG-12-GFP-AID fluorescent signal. Example nuclei are circled with dashed lines. Scale bars: 5 µm. **B.** Violin plots with overlaid notched boxplots of distributions of integrated intensities of transcription dots in the different conditions. Data normalized to the control strain, DLG-1::EGFP::mIAA7 and treated for 30 minutes with DMSO. Data for DLG-1(-) and DLG-1(+) are the same as in Fig. 5C coming from the same set of experiments as a reference. *** = p < 0.001. For raw data and statistics, see Table S4. **C,D,E.** Histograms of distributions of fold values across strains and conditions from (B) and Fig. 5C,D. Fold values are on the x axis and binned into 30 intervals, displayed in separate panels for each strain. Bin counts are shown on the y axis, allowing comparison of distribution shapes within each condition.

## Supplementary table legends

**Table S1.** List of *C. elegans* strains and bacterial RNAi clones used in this study. Sheet1: list of *C. elegans* strains showing strain name, genotype, and reference. Sheet2: list of bacterial RNAi clones showing target gene to downregulate, which library they come from, plasmid name for custom-made clones, and reference. https://doi.org/10.5281/zenodo.21513080

**Table S2.** List of smFISH probes. List of the eighteen probes designed on *ajm-1* intron sequences, containing a FlpX arm. https://doi.org/10.5281/zenodo.21513080

**Table S3.** Raw data for quantitation on live imaging. Each sheet contains the list of data corresponding to the figure panel assessed on the sheet name. Columns show: ID (*i.e*., short name for the strains used – same as in corresponding graphs), min (*i.e*., background), max (*i.e*., actual signal), BG_removal (*i.e*., max – min), AverageREF (*i.e*., average of the reference strain the data have been normalized to), and Norm_intensity (*i.e*., BG_removal / AverageREF). https://doi.org/10.5281/zenodo.21513080

**Table S4.** Raw data for quantitation on smFISH imaging. Each sheet contains the list of data corresponding to the figure panel assessed on the sheet name. https://doi.org/10.5281/zenodo.21513080 Columns show:

| Column Name | Description |
| --- | --- |
| image_name | File name of the original image from which the dot was detected. |
| image_index | Numerical index identifying the image within the dataset. |
| image_min | Minimum pixel intensity value in the image. |
| image_max | Maximum pixel intensity value in the image. |
| image_mean | Mean pixel intensity of the image. |
| image_std | SD of pixel intensities in the image. |
| image_median | Median pixel intensity value of the image. |
| image_mode | Most frequently occurring pixel intensity value in the image. |
| **description** | Describes the channel of origin ('dot1' for single channel, or 'dot1' and 'dot2' for dual channel). |
| **object_id** | Unique identifier assigned to each segmented dot within an image. |
| **pixel_volume** | Total pixel/voxel count for given dot segmentation mask. |
| **volume** | Mesh volume of given dot. |
| **centroid** | Spatial coordinates (Z, Y, X) of the dot's geometric center. |
| **radius_ratio** | Ratio between minimum and maximum radial distances from the centroid. |
| **surface_area** | Mesh surface area of the segmented dot. |
| **sphericity** | Measure of how closely the object shape approximates a perfect sphere. |
| **min_intensity** | Minimum pixel intensity value within the dot mask. |
| **max_intensity** | Maximum pixel intensity value within the dot mask. |
| **mean_intensity** | Mean pixel intensity within the dot mask. |
| **median_intensity** | Median pixel intensity within the dot mask. |
| **std_intensity** | SD of pixel intensities within the dot mask. |
| **background_mean_intensity** | Mean background intensity within the nucleus crop, excluding dot mask pixels. |
| **condition** | Experimental condition under which the sample was acquired. |
| **treatment_group** | Assigned treatment or control group (CTR used as normalization reference). |
| **rescaled_intensity** | $\text{mean\_intensity} - \text{background\_mean\_intensity}$ |
| **integrated_intensity** | $\text{volume} * \text{rescaled\_intensity}$ |
| **volume_normalizer** | Reference value used to normalize dot volume (mean volume of CTR group). |
| **intensity_normalizer** | Reference value used to normalize dot intensity (mean integrated intensity of CTR group). |
| **normalized_volume** | $\text{volume} / \text{volume\_normalizer}$ |
| **normalized_intensity** | $\text{integrated\_intensity} / \text{intensity\_normalizer}$ |

**Table S5.** List of reagents and results of the RT-qPCR. Sheet1: primer list for the tested genes (exonic and intronic). Sheet2: results summary of the data plotted in Fig. S7E. Sheet3: Raw data. https://doi.org/10.5281/zenodo.21513080

## References

1. Campbell, H.K., Maiers, J.L., and DeMali, K.A. (2017). Interplay between tight junctions & adherens junctions. Experimental Cell Research 358, 39–44. 10.1016/j.yexcr.2017.03.061.

2. Campàs, O., Noordstra, I., and Yap, A.S. (2024). Adherens junctions as molecular regulators of emergent tissue mechanics. Nat Rev Mol Cell Biol 25, 252–269. 10.1038/s41580-023-00688-7.

3. Harris, T.J.C., and Tepass, U. (2010). Adherens junctions: from molecules to morphogenesis. Nat Rev Mol Cell Biol 11, 502–514. 10.1038/nrm2927.

4. Hartsock, A., and Nelson, W.J. (2008). Adherens and tight junctions: Structure, function and connections to the actin cytoskeleton. Biochimica et Biophysica Acta (BBA) - Biomembranes 1778, 660–669. 10.1016/j.bbamem.2007.07.012.

5. Liu, J., Li, J., Ren, Y., and Liu, P. (2014). DLG5 in Cell Polarity Maintenance and Cancer Development. Int. J. Biol. Sci. 10, 543–549. 10.7150/ijbs.8888.

6. Parrish, A.R. (2017). The impact of aging on epithelial barriers. Tissue Barriers 5, e1343172. 10.1080/21688370.2017.1343172.

7. Yu, Y., and Elble, R. (2016). Homeostatic Signaling by Cell–Cell Junctions and Its Dysregulation during Cancer Progression. JCM 5, 26. 10.3390/jcm5020026.

8. Carvalho, C.A., and Broday, L. (2020). Game of Tissues: How the Epidermis Thrones C. elegans Shape. JDB 8, 7. 10.3390/jdb8010007.

9. Czerniak, N.D., Dierkes, K., D’Angelo, A., Colombelli, J., and Solon, J. (2016). Patterned Contractile Forces Promote Epidermal Spreading and Regulate Segment Positioning during Drosophila Head Involution. Current Biology 26, 1895–1901. 10.1016/j.cub.2016.05.027.

10. Takeichi, M. (2014). Dynamic contacts: rearranging adherens junctions to drive epithelial remodelling. Nat Rev Mol Cell Biol 15, 397–410. 10.1038/nrm3802.

11. Rosenblatt, J., Raff, M.C., and Cramer, L.P. (2001). An epithelial cell destined for apoptosis signals its neighbors to extrude it by an actin- and myosin-dependent mechanism. Current Biology 11, 1847–1857. 10.1016/S0960-9822(01)00587-5.

12. Fu, Y., Li, X., Fan, B., Zhu, C., and Chen, Z. (2022). Chloroplasts Protein Quality Control and Turnover: A Multitude of Mechanisms. IJMS 23, 7760. 10.3390/ijms23147760.

13. Christianson, J.C., Jarosch, E., and Sommer, T. (2023). Mechanisms of substrate processing during ER-associated protein degradation. Nat Rev Mol Cell Biol 24, 777–796. 10.1038/s41580-023-00633-8.

14. Buckley, C.E., and St Johnston, D. (2022). Apical–basal polarity and the control of epithelial form and function. Nat Rev Mol Cell Biol 23, 559–577. 10.1038/s41580-022-00465-y.

15. Lynch, A., M. (2009). The assembly and maintenance of epithelial junctions in C. elegans. Front Biosci Volume, 1414. 10.2741/3316.

16. Mira-Osuna, M., and Borgne, R.L. (2024). Assembly, dynamics and remodeling of epithelial cell junctions throughout development. Development 151, dev201086. 10.1242/dev.201086.

17. Cordova-Burgos, L., Patel, F.B., and Soto, M.C. (2021). E-Cadherin/HMR-1 Membrane Enrichment Is Polarized by WAVE-Dependent Branched Actin. JDB 9, 19. 10.3390/jdb9020019.

18. Hardin, J., Lynch, A., Loveless, T., and Pettitt, J. (2013). Cadherins and Their Partners in the Nematode Worm Caenorhabditis elegans. In Progress in Molecular Biology and Translational Science (Elsevier), pp. 239–262. 10.1016/B978-0-12-394311-8.00011-X.

19. Lockwood, C., Zaidel-Bar, R., and Hardin, J. (2008). The C. elegans Zonula Occludens Ortholog Cooperates with the Cadherin Complex to Recruit Actin during Morphogenesis. Current Biology 18, 1333–1337. 10.1016/j.cub.2008.07.086.

20. Jud, M.C., Lowry, J., Padilla, T., Clifford, E., Yang, Y., Fennell, F., Miller, A.K., Hamill, D., Harvey, A.M., Avila-Zavala, M., et al. (2021). A genetic screen for temperature-sensitive morphogenesis-defective *Caenorhabditis elegans* mutants. G3 Genes|Genomes|Genetics 11, jkab026. 10.1093/g3journal/jkab026.

21. Costa, M., Raich, W., Agbunag, C., Leung, B., Hardin, J., and Priess, J.R. (1998). A Putative Catenin–Cadherin System Mediates Morphogenesis of the *Caenorhabditis elegans* Embryo. The Journal of Cell Biology 141, 297–308. 10.1083/jcb.141.1.297.

22. Tocchini, C., Rohner, M., Guerard, L., Ray, P., Von Stetina, S.E., and Mango, S.E. (2021). Translation-dependent mRNA localization to *Caenorhabditis elegans* adherens junctions. Development 148, dev200027. 10.1242/dev.200027.

23. Tocchini, C., and Mango, S.E. (2024). An adapted MS2-MCP system to visualize endogenous cytoplasmic mRNA with live imaging in Caenorhabditis elegans. PLoS Biol 22, e3002526. 10.1371/journal.pbio.3002526.

24. Lautier, O., Penzo, A., Rouvière, J.O., Chevreux, G., Collet, L., Loïodice, I., Taddei, A., Devaux, F., Collart, M.A., and Palancade, B. (2021). Co-translational assembly and localized translation of nucleoporins in nuclear pore complex biogenesis. Molecular Cell 81, 2417–2427.e5. 10.1016/j.molcel.2021.03.030.

25. Van Der Salm, E., Koelewijn, E., Schroeder, M., Van Der Maas, E., Jarosińska, O., Eeken, M., and Ruijtenberg, S. (2025). Measuring and manipulating localized translation of *erm-1* in the *C. elegans* embryo. Development 152, dev204435. 10.1242/dev.204435.

26. Galy, V., Mattaj, I.W., and Askjaer, P. Caenorhabditis elegans Nucleoporins Nup93 and Nup205 Determine the Limit of Nuclear Pore Complex Size Exclusion In Vivo□D V □.

27. Wu, J., Matunis, M.J., Kraemer, D., Blobel, G., and Coutavas, E. (1995). Nup358, a Cytoplasmically Exposed Nucleoporin with Peptide Repeats, Ran-GTP Binding Sites, Zinc Fingers, a Cyclophilin A Homologous Domain, and a Leucine-rich Region. Journal of Biological Chemistry 270, 14209–14213. 10.1074/jbc.270.23.14209.

28. Heppert, J.K., Pani, A.M., Roberts, A.M., Dickinson, D.J., and Goldstein, B. (2018). A CRISPR Tagging-Based Screen Reveals Localized Players in Wnt-Directed Asymmetric Cell Division. Genetics 208, 1147–1164. 10.1534/genetics.117.300487.

29. Lee, C., Shin, H., and Kimble, J. (2019). Dynamics of Notch-Dependent Transcriptional Bursting in Its Native Context. Developmental Cell 50, 426–435.e4. 10.1016/j.devcel.2019.07.001.

30. Negishi, T., Kitagawa, S., Horii, N., Tanaka, Y., Haruta, N., Sugimoto, A., Sawa, H., Hayashi, K., Harata, M., and Kanemaki, M.T. (2022). The auxin-inducible degron 2 (AID2) system enables controlled protein knockdown during embryogenesis and development in *Caenorhabditis elegans*. Genetics 220, iyab218. 10.1093/genetics/iyab218.

31. Sepers, J.J., Verstappen, N.H.M., Vo, A.A., Ragle, J.M., Ruijtenberg, S., Ward, J.D., and Boxem, M. (2022). The mIAA7 degron improves auxin-mediated degradation in *Caenorhabditis elegans*. G3 Genes|Genomes|Genetics 12, jkac222. 10.1093/g3journal/jkac222.

32. McMahon, L., Legouis, R., Vonesch, J.-L., and Labouesse, M. (2001). Assembly of *C. elegans* apical junctions involves positioning and compaction by LET-413 and protein aggregation by the MAGUK protein DLG-1. Journal of Cell Science 114, 2265–2277. 10.1242/jcs.114.12.2265.

33. El-Brolosy, M.A., Kontarakis, Z., Rossi, A., Kuenne, C., Günther, S., Fukuda, N., Kikhi, K., Boezio, G.L.M., Takacs, C.M., Lai, S.-L., et al. (2019). Genetic compensation triggered by mutant mRNA degradation. Nature 568, 193–197. 10.1038/s41586-019-1064-z.

34. Serobyan, V., Kontarakis, Z., El-Brolosy, M.A., Welker, J.M., Tolstenkov, O., Saadeldein, A.M., Retzer, N., Gottschalk, A., Wehman, A.M., and Stainier, D.Y. (2020). Transcriptional adaptation in Caenorhabditis elegans. eLife 9, e50014. 10.7554/eLife.50014.

35. Falcucci, L., Juvik, B., and Stainier, D.Y. (2025). Transcriptional adaptation: where mRNA decay meets genetic compensation. Current Opinion in Genetics & Development 93, 102369. 10.1016/j.gde.2025.102369.

36. Pickles, S., Vigié, P., and Youle, R.J. (2018). Mitophagy and Quality Control Mechanisms in Mitochondrial Maintenance. Current Biology 28, R170–R185. 10.1016/j.cub.2018.01.004.

37. Brangwynne, C.P., MacKintosh, F.C., Kumar, S., Geisse, N.A., Talbot, J., Mahadevan, L., Parker, K.K., Ingber, D.E., and Weitz, D.A. (2006). Microtubules can bear enhanced compressive loads in living cells because of lateral reinforcement. The Journal of Cell Biology 173, 733–741. 10.1083/jcb.200601060.

38. Rey-Suarez, I., Rogers, N., Kerr, S., Shroff, H., and Upadhyaya, A. (2021). Actomyosin dynamics modulate microtubule deformation and growth during T-cell activation. MBoC 32, 1641–1653. 10.1091/mbc.E20-10-0685.

39. Crisp, M., Liu, Q., Roux, K., Rattner, J.B., Shanahan, C., Burke, B., Stahl, P.D., and Hodzic, D. (2006). Coupling of the nucleus and cytoplasm: Role of the LINC complex. The Journal of Cell Biology 172, 41–53. 10.1083/jcb.200509124.

40. Janota, C.S., Calero-Cuenca, F.J., and Gomes, E.R. (2020). The role of the cell nucleus in mechanotransduction. Current Opinion in Cell Biology 63, 204–211. 10.1016/j.ceb.2020.03.001.

41. Tzur, Y.B., Wilson, K.L., and Gruenbaum, Y. (2006). SUN-domain proteins: “Velcro” that links the nucleoskeleton to the cytoskeleton. Nat Rev Mol Cell Biol 7, 782–788. 10.1038/nrm2003.

42. Worman, H.J., and Gundersen, G.G. (2006). Here come the SUNs: a nucleocytoskeletal missing link. Trends in Cell Biology 16, 67–69. 10.1016/j.tcb.2005.12.006.

43. Rothballer, A., and Kutay, U. (2013). The diverse functional LINCs of the nuclear envelope to the cytoskeleton and chromatin. Chromosoma 122, 415–429. 10.1007/s00412-013-0417-x.

44. Zhou, K., and Hanna-Rose, W. (2010). Movers and shakers or anchored: *Caenorhabditis elegans* nuclei achieve it with KASH/SUN. Developmental Dynamics 239, 1352–1364. 10.1002/dvdy.22226.

45. Cohen-Fix, O., and Askjaer, P. (2017). Cell Biology of the *Caenorhabditis elegans* Nucleus. Genetics 205, 25–59. 10.1534/genetics.116.197160.

46. Lawrence, K.S., Tapley, E.C., Cruz, V.E., Li, Q., Aung, K., Hart, K.C., Schwartz, T.U., Starr, D.A., and Engebrecht, J. (2016). LINC complexes promote homologous recombination in part through inhibition of nonhomologous end joining. Journal of Cell Biology 215, 801–821. 10.1083/jcb.201604112.

47. Starr, D.A. (2019). A network of nuclear envelope proteins and cytoskeletal force generators mediates movements of and within nuclei throughout *Caenorhabditis elegans* development. Exp Biol Med (Maywood) 244, 1323–1332. 10.1177/1535370219871965.

48. Assémat, E., Bazellières, E., Pallesi-Pocachard, E., Le Bivic, A., and Massey-Harroche, D. (2008). Polarity complex proteins. Biochimica et Biophysica Acta (BBA) – Biomembranes 1778, 614–630. 10.1016/j.bbamem.2007.08.029.

49. Won, S., Levy, J.M., Nicoll, R.A., and Roche, K.W. (2017). MAGUKs: multifaceted synaptic organizers. Current Opinion in Neurobiology 43, 94–101. 10.1016/j.conb.2017.01.006.

50. Zanin-Zhorov, A., Lin, J., Scher, J., Kumari, S., Blair, D., Hippen, K.L., Blazar, B.R., Abramson, S.B., Lafaille, J.J., and Dustin, M.L. (2012). Scaffold protein Disc large homolog 1 is required for T-cell receptor-induced activation of regulatory T-cell function. Proc. Natl. Acad. Sci. U.S.A. 109, 1625–1630. 10.1073/pnas.1110120109.

51. Xavier, R., Rabizadeh, S., Ishiguro, K., Andre, N., Ortiz, J.B., Wachtel, H., Morris, D.G., Lopez-Ilasaca, M., Shaw, A.C., Swat, W., et al. (2004). Discs large (Dlg1) complexes in lymphocyte activation. The Journal of Cell Biology 166, 173–178. 10.1083/jcb.200309044.

52. Cavatorta, A.L., Di Gregorio, A., Bugnon Valdano, M., Marziali, F., Cabral, M., Bottai, H., Cittadini, J., Nocito, A.L., and Gardiol, D. (2017). DLG1 polarity protein expression associates with the disease progress of low-grade cervical intraepithelial lesions. Experimental and Molecular Pathology 102, 65–69. 10.1016/j.yexmp.2016.12.008.

53. Zhu, G.-D., OuYang, S., Liu, F., Zhu, Z.-G., Jiang, F.-N., and Zhang, B. (2017). Elevated Expression of DLG1 Is Associated with Poor Prognosis in Patients with Colorectal Cancer. 47.

54. Lécuyer, E., Yoshida, H., Parthasarathy, N., Alm, C., Babak, T., Cerovina, T., Hughes, T.R., Tomancak, P., and Krause, H.M. (2007). Global Analysis of mRNA Localization Reveals a Prominent Role in Organizing Cellular Architecture and Function. Cell 131, 174–187. 10.1016/j.cell.2007.08.003.

55. Wan, Y., El Kholtei, J., Jenie, I., Colomer-Rosell, M., Liu, J., Acedo, J.N., Du, L.Y., Codina-Tobias, M., Wang, M., Sawh, A., et al. (2024). Whole-embryo Spatial Transcriptomics at Subcellular Resolution from Gastrulation to Organogenesis. Preprint at Developmental Biology, https://doi.org/10.1101/2024.08.27.609868 10.1101/2024.08.27.609868.

56. Costa, G., Bradbury, J.J., Tarannum, N., and Herbert, S.P. (2020). RAB13 mRNA compartmentalisation spatially orients tissue morphogenesis. EMBO J 39, EMBJ2020106003. 10.15252/embj.2020106003.

57. Norris, M.L., and Mendell, J.T. (2023). Localization of *Kif1c* mRNA to cell protrusions dictates binding partner specificity of the encoded protein. Genes Dev. 37, 191–203. 10.1101/gad.350320.122.

58. Brendza, R.P., Serbus, L.R., Duffy, J.B., and Saxton, W.M. (2000). A Function for Kinesin I in the Posterior Transport of *oskar* mRNA and Staufen Protein. Science 289, 2120–2122. 10.1126/science.289.5487.2120.

59. St Johnston, D. (1995). The intracellular localization of messenger RNAs. Cell 81, 161–170. 10.1016/0092-8674(95)90324-0.

60. Micklem, D.R. (1995). mRNA Localisation during Development. Developmental Biology 172, 377–395. 10.1006/dbio.1995.8048.

61. Das, S., Vera, M., Gandin, V., Singer, R.H., and Tutucci, E. (2021). Intracellular mRNA transport and localized translation. Nat Rev Mol Cell Biol 22, 483–504. 10.1038/s41580-021-00356-8.

62. Fazal, F.M., Han, S., Parker, K.R., Kaewsapsak, P., Xu, J., Boettiger, A.N., Chang, H.Y., and Ting, A.Y. (2019). Atlas of Subcellular RNA Localization Revealed by APEX-Seq. Cell 178, 473–490.e26. 10.1016/j.cell.2019.05.027.

63. Mofatteh, M., 1 Lincoln College, University of Oxford, Turl Street, Oxford, OX1 3DR, United Kingdom, 2 Merton College, University of Oxford, Merton Street, Oxford, OX1 4DJ, United Kingdom, and 3 Sir William Dunn School of Pathology, University of Oxford, South Parks Road, Oxford, OX1 3RE, United Kingdom (2020). mRNA localization and local translation in neurons. AIMS Neuroscience 7, 299–310. 10.3934/Neuroscience.2020016.

64. Turner-Bridger, B., Caterino, C., and Cioni, J.-M. (2020). Molecular mechanisms behind mRNA localization in axons. Open Biol. 10, 200177. 10.1098/rsob.200177.

65. Goering, R., Arora, A., Pockalny, M.C., and Taliaferro, J.M. (2023). RNA localization mechanisms transcend cell morphology. eLife 12, e80040. 10.7554/eLife.80040.

66. Mason, D.E., Madsen, T.D., Gasparski, A.N., Jiwnani, N., Lechler, T., Weigert, R., Iglesias-Bartolome, R., and Mili, S. (2024). Control of Epithelial Tissue Organization by mRNA Localization. Preprint at Cell Biology, https://doi.org/10.1101/2024.12.02.626432 10.1101/2024.12.02.626432.

67. Novoselsky, R., Golani, O., Barkai, T., Kedmi, M., Goliand, I., Fine, M., Kent, I., Nachmany, I., and Itzkovitz, S. (2026). Subcellular mRNA localization patterns across tissues resolved with spatial transcriptomics. Nat Commun 17, 5466. 10.1038/s41467-026-72156-7.

68. Moor, A.E., Golan, M., Massasa, E.E., Lemze, D., Weizman, T., Shenhav, R., Baydatch, S., Mizrahi, O., Winkler, R., Golani, O., et al. (2017). Global mRNA polarization regulates translation efficiency in the intestinal epithelium. Science 357, 1299–1303. 10.1126/science.aan2399.

69. Arceo, X.G., Koslover, E.F., Zid, B.M., and Brown, A.I. (2022). Mitochondrial mRNA localization is governed by translation kinetics and spatial transport. PLoS Comput Biol 18, e1010413. 10.1371/journal.pcbi.1010413.

70. Voigt, F., Zhang, H., Cui, X.A., Triebold, D., Liu, A.X., Eglinger, J., Lee, E.S., Chao, J.A., and Palazzo, A.F. (2017). Single-Molecule Quantification of Translation-Dependent Association of mRNAs with the Endoplasmic Reticulum. Cell Reports 21, 3740–3753. 10.1016/j.celrep.2017.12.008.

71. Semotok, J.L., Cooperstock, R.L., Pinder, B.D., Vari, H.K., Lipshitz, H.D., and Smibert, C.A. (2005). Smaug Recruits the CCR4/POP2/NOT Deadenylase Complex to Trigger Maternal Transcript Localization in the Early Drosophila Embryo. Current Biology 15, 284–294. 10.1016/j.cub.2005.01.048.

72. Ding, D., Parkhurst, S.M., and Lipshitz, H.D. (1993). Different genetic requirements for anterior RNA localization revealed by the distribution of Adducin-like transcripts during Drosophila oogenesis. Proc. Natl. Acad. Sci. U.S.A. 90, 2512–2516. 10.1073/pnas.90.6.2512.

73. Mendonsa, S., Von Kügelgen, N., Dantsuji, S., Ron, M., Breimann, L., Baranovskii, A., Lödige, I., Kirchner, M., Fischer, M., Zerna, N., et al. (2023). Massively parallel identification of mRNA localization elements in primary cortical neurons. Nat Neurosci. 10.1038/s41593-022-01243-x.

74. Poulopoulos, A., Murphy, A.J., Ozkan, A., Davis, P., Hatch, J., Kirchner, R., and Macklis, J.D. (2019). Subcellular transcriptomes and proteomes of developing axon projections in the cerebral cortex. Nature 565, 356–360. 10.1038/s41586-018-0847-y.

75. Yergert, K.M., Doll, C.A., O’Rouke, R., Hines, J.H., and Appel, B. (2021). Identification of 3′ UTR motifs required for mRNA localization to myelin sheaths in vivo. PLoS Biol 19, e3001053. 10.1371/journal.pbio.3001053.

76. Balchin, D., Hayer-Hartl, M., and Hartl, F.U. (2016). In vivo aspects of protein folding and quality control. Science 353, aac4354. 10.1126/science.aac4354.

77. Song, J., Herrmann, J.M., and Becker, T. (2021). Quality control of the mitochondrial proteome. Nat Rev Mol Cell Biol 22, 54–70. 10.1038/s41580-020-00300-2.

78. Wiseman, R.L., Mesgarzadeh, J.S., and Hendershot, L.M. (2022). Reshaping endoplasmic reticulum quality control through the unfolded protein response. Molecular Cell 82, 1477–1491. 10.1016/j.molcel.2022.03.025.

79. Rodriguez-Rocha, H., Garcia-Garcia, A., Panayiotidis, M.I., and Franco, R. (2011). DNA damage and autophagy. Mutation Research/Fundamental and Molecular Mechanisms of Mutagenesis 711, 158–166. 10.1016/j.mrfmmm.2011.03.007.

80. Lejeune, F. (2022). Nonsense-Mediated mRNA Decay, a Finely Regulated Mechanism. Biomedicines 10, 141. 10.3390/biomedicines10010141.

81. Pontisso, I., Ornelas-Guevara, R., Combettes, L., and Dupont, G. (2023). A journey in UPR modelling. Biology of the Cell 115, 2200111. 10.1111/boc.202200111.

82. Elborn, J.S. (2016). Cystic fibrosis. The Lancet 388, 2519–2531. 10.1016/S0140-6736(16)00576-6.

83. Hetz, C., and Papa, F.R. (2018). The Unfolded Protein Response and Cell Fate Control. Molecular Cell 69, 169–181. 10.1016/j.molcel.2017.06.017.

84. Hipp, M.S., Kasturi, P., and Hartl, F.U. (2019). The proteostasis network and its decline in ageing. Nat Rev Mol Cell Biol 20, 421–435. 10.1038/s41580-019-0101-y.

85. Liu, B.-H., Xu, C.-Z., Liu, Y., Lu, Z.-L., Fu, T.-L., Li, G.-R., Deng, Y., Luo, G.-Q., Ding, S., Li, N., et al. (2024). Mitochondrial quality control in human health and disease. Military Med Res 11, 32. 10.1186/s40779-024-00536-5.

86. Mahat, D.B., and Lis, J.T. (2017). Use of conditioned media is critical for studies of regulation in response to rapid heat shock. Cell Stress and Chaperones 22, 155–162. 10.1007/s12192-016-0737-x.

87. Dupont, S., and Wickström, S.A. (2022). Mechanical regulation of chromatin and transcription. Nat Rev Genet 23, 624–643. 10.1038/s41576-022-00493-6.

88. Yonemura, S., Wada, Y., Watanabe, T., Nagafuchi, A., and Shibata, M. (2010). α-Catenin as a tension transducer that induces adherens junction development. Nat Cell Biol 12, 533– 542. 10.1038/ncb2055.

89. Le Duc, Q., Shi, Q., Blonk, I., Sonnenberg, A., Wang, N., Leckband, D., and De Rooij, J. (2010). Vinculin potentiates E-cadherin mechanosensing and is recruited to actin-anchored sites within adherens junctions in a myosin II–dependent manner. Journal of Cell Biology 189, 1107–1115. 10.1083/jcb.201001149.

90. Vuong-Brender, T.T.K., Suman, S.K., and Labouesse, M. (2017). The apical ECM preserves embryonic integrity and distributes mechanical stress during morphogenesis. Development, dev.150383. 10.1242/dev.150383.

91. Portnoy, V., Huang, V., Place, R.F., and Li, L. (2011). Small RNA and transcriptional upregulation. WIREs RNA 2, 748–760. 10.1002/wrna.90.

92. Seroussi, U., Li, C., Sundby, A.E., Lee, T.L., Claycomb, J.M., and Saltzman, A.L. (2022). Mechanisms of epigenetic regulation by C. elegans nuclear RNA interference pathways. Seminars in Cell & Developmental Biology 127, 142–154. 10.1016/j.semcdb.2021.11.018.

93. Malone, C.J., Fixsen, W.D., Horvitz, H.R., and Han, M. (1999). UNC-84 localizes to the nuclear envelope and is required for nuclear migration and anchoring during *C. elegans* development. Development 126, 3171–3181. 10.1242/dev.126.14.3171.

94. Starr, D.A., Hermann, G.J., Malone, C.J., Fixsen, W., Priess, J.R., Horvitz, H.R., and Han, M. (2001). *unc-83* encodes a novel component of the nuclear envelope and is essential for proper nuclear migration. Development 128, 5039–5050. 10.1242/dev.128.24.5039.

95. Starr, D.A., and Han, M. (2002). Role of ANC-1 in Tethering Nuclei to the Actin Cytoskeleton. Science 298, 406–409. 10.1126/science.1075119.

96. Jahed, Z., Domkam, N., Ornowski, J., Yerima, G., and Mofrad, M.R.K. (2021). Molecular models of LINC complex assembly at the nuclear envelope. Journal of Cell Science 134, jcs258194. 10.1242/jcs.258194.

97. Purushothaman, D., Bianchi, L.F., Penkov, D., Poli, A., Li, Q., Vermezovic, J., Pramotton, F.M., Choudhary, R., Pennacchio, F.A., Sommariva, E., et al. (2022). The transcription factor PREP1(PKNOX1) regulates nuclear stiffness, the expression of LINC complex proteins and mechanotransduction. Commun Biol 5, 456. 10.1038/s42003-022-03406-9.

98. Garner, K.E.L., Salter, A., Lau, C.K., Gurusaran, M., Villemant, C.M., Granger, E.P., McNee, G., Woodman, P.G., Davies, O.R., Burke, B.E., et al. (2023). The meiotic LINC complex component KASH5 is an activating adaptor for cytoplasmic dynein. Journal of Cell Biology 222, e202204042. 10.1083/jcb.202204042.

99. Carvalho, C., Moreira, M., Barbosa, D.J., Chan, F.-Y., Koehnen, C.B., Teixeira, V., Rocha, H., Green, M., Carvalho, A.X., Cheerambathur, D.K., et al. (2025). ZYG-12/Hook’s dual role as a dynein adaptor for early endosomes and nuclei is regulated by alternative splicing of its cargo binding domain. MBoC 36, ar19. 10.1091/mbc.E24-08-0364.

100. Mlynarczyk-Evans, S., and Villeneuve, A.M. (2017). Time-Course Analysis of Early Meiotic Prophase Events Informs Mechanisms of Homolog Pairing and Synapsis in *Caenorhabditis elegans*. Genetics 207, 103–114. 10.1534/genetics.117.204172.

101. Penkner, A.M., Fridkin, A., Gloggnitzer, J., Baudrimont, A., Machacek, T., Woglar, A., Csaszar, E., Pasierbek, P., Ammerer, G., Gruenbaum, Y., et al. (2009). Meiotic Chromosome Homology Search Involves Modifications of the Nuclear Envelope Protein Matefin/SUN-1. Cell 139, 920–933. 10.1016/j.cell.2009.10.045.

102. Sato, A., Isaac, B., Phillips, C.M., Rillo, R., Carlton, P.M., Wynne, D.J., Kasad, R.A., and Dernburg, A.F. (2009). Cytoskeletal Forces Span the Nuclear Envelope to Coordinate Meiotic Chromosome Pairing and Synapsis. Cell 139, 907–919. 10.1016/j.cell.2009.10.039.

103. Malone, C.J., Misner, L., Le Bot, N., Tsai, M.-C., Campbell, J.M., Ahringer, J., and White, J.G. (2003). The C. elegans Hook Protein, ZYG-12, Mediates the Essential Attachment between the Centrosome and Nucleus. Cell 115, 825–836. 10.1016/S0092-8674(03)00985-1.

104. Dogterom, M., and Koenderink, G.H. (2019). Actin–microtubule crosstalk in cell biology. Nat Rev Mol Cell Biol 20, 38–54. 10.1038/s41580-018-0067-1.

105. Hardin, J. (2005). Epidermal morphogenesis. WormBook. 10.1895/wormbook.1.35.1.

106. Han, M., Fu, M.L., Zhu, Y., Choi, A.A., Li, E., Bezney, J., Cai, S., Miles, L., Ma, Y., and Qi, L.S. (2025). Programmable control of spatial transcriptome in live cells and neurons. Nature 643, 241–251. 10.1038/s41586-025-09020-z.

107. Kamath, R.S., Fraser, A.G., Dong, Y., Poulin, G., Durbin, R., Gotta, M., Kanapin, A., Le Bot, N., Moreno, S., Sohrmann, M., et al. (2003). Systematic functional analysis of the Caenorhabditis elegans genome using RNAi. Nature 421, 231–237. 10.1038/nature01278.

108. Imbert, A., Ouyang, W., Safieddine, A., Coleno, E., Zimmer, C., Bertrand, E., Walter, T., and Mueller, F. (2022). FISH-quant v2: a scalable and modular tool for smFISH image analysis. RNA 28, 786–795. 10.1261/rna.079073.121.

109. Frøkjær-Jensen, C., Wayne Davis, M., Hopkins, C.E., Newman, B.J., Thummel, J.M., Olesen, S.-P., Grunnet, M., and Jorgensen, E.M. (2008). Single-copy insertion of transgenes in Caenorhabditis elegans. Nat Genet 40, 1375–1383. 10.1038/ng.248.

110. Mueller, F., Senecal, A., Tantale, K., Marie-Nelly, H., Ly, N., Collin, O., Basyuk, E., Bertrand, E., Darzacq, X., and Zimmer, C. (2013). FISH-quant: automatic counting of transcripts in 3D FISH images. Nat Methods 10, 277–278. 10.1038/nmeth.2406.

111. Ghanta, K.S., Ishidate, T., and Mello, C.C. (2021). Microinjection for precision genome editing in Caenorhabditis elegans. STAR Protocols 2, 100748. 10.1016/j.xpro.2021.100748.

112. Tsanov, N., Samacoits, A., Chouaib, R., Traboulsi, A.-M., Gostan, T., Weber, C., Zimmer, C., Zibara, K., Walter, T., Peter, M., et al. (2016). smiFISH and FISH-quant – a flexible single RNA detection approach with super-resolution capability. Nucleic Acids Res 44, e165–e165. 10.1093/nar/gkw784.

113. Francis, R., and Waterston, R.H. (1991). Muscle cell attachment in Caenorhabditis elegans. The Journal of cell biology 114, 465–479. 10.1083/jcb.114.3.465.

